# Reliable Detection of SGLT2 Protein through Knockout-based Antibody Characterization and Optimized Procedures

**DOI:** 10.1101/2025.08.31.672288

**Authors:** Takuo Hirose, Hiroki Ito, Akari Endo, Shigemitsu Sato, Chika Takahashi, Takahito Kaburagi, Kentaro Yano, Risa Ishikawa, Ayaka Kamada, Ikuko Oba-Yabana, Michihiro Satoh, Kento Morozumi, Yasuhiro Kaiho, Yasuhiro Nakamura, Keiju Kamijo, Wako Yumura, Takefumi Mori

## Abstract

**Introduction:** Sodium-glucose cotransporter 2 (SGLT2) is a key mediator of renal glucose reabsorption. Its pharmacological inhibition exerts cardio- and reno-protective benefits. Despite widespread clinical interest, reliable detection of SGLT2 protein remains challenging due to concerns regarding the specificity of available antibodies.

**Methods:** This study assessed the specificity of eight commercially available anti-SGLT2 antibodies by immunohistochemistry and Western blotting. Genetically engineered *Sglt2*-deficient mice and rats, generated via clustered regularly interspaced short palindromic repeats/CRISPR-associated protein 9 (CRISPR/Cas9) technology, were employed as definitive negative controls. Additionally, human kidney tissues, including renal cell carcinoma samples, were analyzed.

**Results:** Among the antibodies tested, few exhibited robust specificity, characterized by substantial immunostaining with minimal background in wild-type kidney tissues and complete absence of staining in *Sglt2*-deficient samples. In renal cell carcinoma samples, a validated antibody detected SGLT2 immunostaining in proximal tubules of non-tumor regions but not in tumor areas. Subcellular localization studies revealed that SGLT2 was enriched within proximal tubular microvilli, partially overlapping with its co-factor PDZK1IP1 (MAP17). LRP2 (megalin) and NHE3 were placed at the microvillar base and did not colocalize with SGLT2. Western blotting identified a specific SGLT2 band at approximately 55 kDa in kidney lysates using several antibodies under optimized procedures. This band was shifted to approximately 45 kDa after enzymatic removal of N-linked glycans. One antibody detected a weak band at the same molecular mass even in kidney lysates from *Sglt2*-deficient rodents.

**Conclusions:** Considerable variability exists in the specificity of commercially available anti-SGLT2 antibodies. Only a limited number of antibodies are suitable for reliable detection of SGLT2 in rodent and human samples. Rigorous antibody characterization, including the use of knockout controls and optimized experimental conditions, is essential to ensure reproducibility and prevent misinterpretation in studies investigating the biological and pathophysiological roles of SGLT2.

## INTRODUCTION

Sodium-glucose cotransporter 2 (SGLT2), encoded by the *solute carrier family 5 member 2* (*SLC5A2*) gene, is predominantly expressed on the apical membrane of the S1 and S2 segments of the renal proximal tubules.^1,2^ SGLT2 plays a central role in renal glucose reabsorption, and its pharmacological inhibition is a cornerstone of type 2 diabetes mellitus management. Beyond glycemic control, SGLT2 inhibitors have demonstrated cardio- and reno-protective effects in clinical studies.^3,4^ Elucidating the mechanisms underlying these benefits requires detailed knowledge of the SGLT2 protein, including tissue distribution, subcellular localization, and post-translational modifications.

While low levels of *SGLT2* mRNA have been reported in several non-renal tissues,^5^ the corresponding protein expression in normal and malignant tissues remains poorly defined. Protein detection techniques, such as immunohistochemistry and Western blotting, depend on specific antigen-antibody interactions. However, concerns regarding antibody specificity, particularly cross-reactivity and non-specific binding, have been raised for numerous targets, including angiotensin II type 1 receptor and CD68.^6–8^ The use of target gene knockout samples is considered a valuable approach for antibody characterization,^8–11^ but a systematic assessment of anti-SGLT2 antibodies remains to be conducted.

Recent advances in genome editing, particularly the clustered regularly interspaced short palindromic repeats/CRISPR-associated protein 9 (CRISPR/Cas9) system, have enabled the rapid generation of genetically modified animals.^12,13^ In the present study, we generated *Sglt2*-deficient rodents and evaluated the specificity of commercially available anti-SGLT2 antibodies through immunohistochemical and Western blot analyses.

## METHODS

### Animals

All animal experiments were conducted in accordance with the National Institutes of Health Guide for the Care and Use of Laboratory Animals. The study protocols received approval from the Tohoku Medical and Pharmaceutical University Animal Experiment Committee (approval numbers: A20015, A21011-cn, A22039-cn, A23033-cn, A24037-cn, and A25007-cn). Transgenic experiments were also reviewed and approved by the Tohoku Medical and Pharmaceutical University Safety Committee for Recombinant DNA Experiments (approval numbers: 2020-20, 2022-28, and 2024-03). Eight-week-old C57BL/6 mice (C57BL/6JJmsSlc) and Sprague-Dawley rats (Slc:SD) were purchased from Japan SLC (Shizuoka, Japan). Animals were housed in a specific pathogen-free facility under controlled environmental conditions (temperature: 22±2°C; humidity: 55±10%; and 12-hour light/dark cycle) with *ad libitum* access to standard chow (CE-2; CLEA Japan, Tokyo, Japan) and tap water.

### Generation of target gene-deficient rodents

Target gene-deficient mice and rats were generated using the improved genome-editing via oviductal nucleic acids delivery (*i*-GONAD) technique, a CRISPR/Cas9-based *in vivo* gene editing approach (Supplementary Figure S1-S4).^12,14–17^ Briefly, pregnant females received intra-oviductal injections of CRISPR/Cas9 components containing Cas9 protein, guide RNA (gRNA), and single-stranded oligodeoxynucleotides (ssODNs) into the ampulla on a temperature-controlled (37°C) surgical table. Electroporation of the oviduct was then performed using a NEPA21 electroporator (NEPAGENE, Chiba, Japan). These procedures were conducted at gestational day 0.7 in mice (n = 5 per target gene) and day 0.75 in rats (n = 2 per target gene) under a triple drug anesthesia cocktail: medetomidine (0.15 mg/kg body weight; Maruishi Pharmaceutical, Osaka, Japan), midazolam (2.0 mg/kg body weight; Astellas Pharma, Tokyo, Japan), and butorphanol (2.5 mg/kg body weight; Meiji Seika Pharma, Tokyo, Japan). gRNAs were designed using CHOPCHOP software (http://chopchop.cbu.uib.no/).^18^ The Cas9 protein, gRNAs, and ssODNs were purchased from Integrated DNA Technologies (Alt-R CRISPR-Cas9 System; Coralville, IA).

Treated females and offspring were maintained under standard housing conditions until weaning. At weaning, ear tissue fragment was collected from offspring for genotyping by Sanger sequencing (Azenta, Burlington, MA). ssODN-mediated knock-in animals were selected and crossed to obtain wild-type and knockout littermates. For *Sglt1*-deficient mice, non-homologous end joining (NHEJ) editing was employed, and edited mice with a 2-base pair deletion were selected.

At 8 weeks of age, littermates from three different mothers were euthanized under isoflurane anesthesia. After a mid-abdominal incision, urine was collected from the urinary bladder for glucosuria testing using dipstick paper (product number E-UR65, Eiken Chemical, Tokyo, Japan). Kidneys and hearts were harvested, and bisected transversely using a disposable surgical blade (product number 750BH2037; KAI CORPORATION, Tokyo, Japan). For histological analysis, tissue fragments were immediately fixed by immersion in 10% neutralized buffered formalin (Mildform 10N, product number 133-10311; FUJIFILM Wako Pure Chemical Corporation, Osaka, Japan) overnight at room temperature on a waving shaker (∼20 rpm, TM-300; AS ONE, Osaka, Japan), processed in an automated tissue processor (Tissue-Tek VIP 6 AI; Sakura Finetek Japan, Tokyo, Japan), and embedded in paraffin. For Western blotting, the kidneys were dissected by visual inspection, snap-frozen in liquid nitrogen, and stored at -80°C until protein extraction for Western blotting.

### Human kidney samples

The use of human samples was carried out in accordance with the Declaration of Helsinki. All protocols involving human tissues were reviewed and approved by the Tohoku Medical and Pharmaceutical University Hospital Institutional Review Board (registration numbers: 2021-2-109 and 2024-2-078-0000).

Human kidney samples were obtained from diagnostic renal biopsies (n = 2), autopsies (n = 8),^19^ and partial nephrectomies for renal cell carcinoma (n = 17, Supplementary Table S1) conducted at Tohoku Medical and Pharmaceutical University Hospital. Paraffin-embedded tissue blocks were prepared by standard procedures in the Division of Diagnostic Pathology of Tohoku Medical and Pharmaceutical University Hospital. Periodic acid-Schiff (PAS) staining was routinely performed at the Histopathology Core Facility of Tohoku Medical and Pharmaceutical University.

### Immunohistochemical staining of anti-SGLT2 antibodies

Immunohistochemical staining was performed as previously described,^19–22^ with minor modifications. Formalin-fixed, paraffin-embedded tissue samples were sliced into 1.5-µm-thick sections for human kidney samples and 4-µm-thick sections for mouse and rat tissue samples by staff at the Histopathology Core Facility of Tohoku Medical and Pharmaceutical University. Sections were mounted onto CREST adhesive glass slides (product number CRE-01; Matsunami Glass Industry, Osaka, Japan). For rodent samples, tissue from wild-type and knockout littermate was placed on the same slide.

After deparaffinization in xylene and rehydration through gradient ethanol series (100%, 100%, 90%, and 80%) and distilled water, heat-induced epitope retrieval was performed using an autoclave (product number SX-500; TOMY, Tokyo, Japan) at 121°C for 5 minutes with either 10 mmol/L citrate buffer (pH 6.0) or 1.0 mmol/L ethylenediaminetetraacetic acid (EDTA) buffer (pH 9.0). Endogenous peroxidase activity was quenched with 0.3% hydrogen peroxide for 10 minutes. Nonspecific binding was blocked using 5% bovine serum albumin (BSA, product number A9647; Sigma-Aldrich, St. Louis, MO)/phosphate buffered saline (PBS, product number T9181; TaKaRa, Osaka, Japan) for 30 minutes at room temperature. Subsequently, the slides were incubated with primary antibodies (Table 1 and Supplementary Table S2) overnight at 4°C in a moist chamber. Eight commercially available anti-SGLT2 antibodies were selected primarily from sources frequently cited in the publications (Table 1), and were diluted in five steps by sequential twofold dilutions using 5% BSA/PBS as the diluent, starting at a concentration higher than that previously reported in the literature or preliminary condition tests conducted in our laboratory. On the next day, slides were washed 3 times with PBS and incubated for 30 minutes at room temperature with a species-specific immuno-peroxidase polymer (Histofine simple stain MAX-PO for human, mouse, or rat; Nichirei Biosciences, Tokyo, Japan) in a moist chamber. After washing 3 times with PBS, the reaction was developed for 2 minutes at room temperature using 3,3’-diaminobenzidine (DAB, product number SK-4100; Vector Laboratories, Newark, CA), followed by three distilled water washes. Counterstaining was performed with Carrazzi’s Hematoxylin Solution (product number 1.15938; Sigma-Aldrich) for 30 sec at room temperature. After dehydration through 4 changes of ethanol (80%, 90%, 100%, and 100%) and cleared in 4 changes of xylene, sections were mounted using NEO micro cover glass (0.13-0.17 mm thickness, product number C024401; Matsunami Glass Industry) and HSR solution (product number 10200; Sysmex, Kobe, Japan). Slides were scanned and digitized using an automatic whole-slide scanner (NanoZoomer-SQ; Hamamatsu Photonics, Hamamatsu, Japan) and the associated software (NDP.view2 ver. 2.9.29; Hamamatsu Photonics).

**Table 1.** Datasheet information of anti-SGLT2 antibodies. ^1^.

| Vendor | Catalog no. | Lot no. | Clone no. | Host species | Conc. (mg/mL) | Antigen | Reactivity species | Application |
| --- | --- | --- | --- | --- | --- | --- | --- | --- |
| Abcam | ab37296 | 1080386-8 | polyclonal | rabbit | 0.5 | mo, N-terminal | hu, mo, rat | WB |
|  | ab85626 | GR3314472-2 | polyclonal | rabbit | 0.9 | hu, 250-350 a.a. | hu<sup>2</sup> | IHC<sup>3,4</sup> |
|  | ab306558 | 1031851-11 | EPR27112-7 | rabbit | 0.52 | proprietary | mo, rat | WB, IHC<sup>4</sup> |
| BiCell Scientific | 20802 | 240139 | polyclonal | rabbit | 0.25 | mo, N-terminal<br>cytoplasmic region 18 a.a. | mo, rat | IF |
| Cell Signaling Technology | 14210 | 2 | polyclonal | rabbit | 0.0815 | hu, near the amino terminus | hu | WB, IP |
| Proteintech | 24654-1-AP | 00111271 | polyclonal | rabbit | 0.3 | hu, 180-311 a.a. | hu, mo, rat | WB, IHC<sup>4</sup>, IF, IP, ELISA |
| Santa Cruz Biotechnology | sc-393350 | A0324 | D-6 | mouse | 0.2 | hu, 220-270 a.a.<sup>5</sup> | hu, mo, rat | WB, IHC, IF, IP, ELISA |
| Sigma-Aldrich | HPA041603 | 000023192 | polyclonal | rabbit | 0.1 | hu, 220-266 a.a. | hu | IHC<sup>4</sup> |

1 On August 31st, 2025, the vendor's webpage was accessed and the printed antibody datasheet supplied at purchase was checked.
2 The product datasheet lists cross-reactivity with mouse, rat, rabbit, cow, dog, pig, and orangutan.
3 The application information on the vendor's webpage (IHC) differs from the printed datasheet supplied with the antibody (WB, IHC, IF), and no datasheet version or update history could be identified.
4 Heat-induced epitope retrieval is recommended using citrate buffer (pH 6.0) for ab85626 and ab306558; TE buffer (pH 9.0) for 24654-1-AP; and retrieval buffer (pH 6.1; Dako, Agilent, Santa Clara, CA, USA) for HPA041603.
5 The antigen region is listed as amino acid residues 226-268 on the product webpage, but as residues 220-270 in the PDF datasheet.
Conc., concentration; hu, human; mo, mouse; a.a., amino acid; WB, Western blotting; IHC, immunohistochemistry; IF, immunofluorescence; IP, immunoprecipitation; ELISA, enzyme-linked immunosorbent assay.

The specificity of the HPA041603 antibody (Sigma-Aldrich) was examined in kidney sections of human biopsy, mouse, and rat by the absorption test. The antibody diluted to working condition (1:2000, 0.05 µg/mL) was preincubated with a 10-fold molar excess of the corresponding antigen (PrEST Antigen SLC5A2, product number APREST82172; Sigma-Aldrich) for 4 hours at 4°C with gentle shaking prior to use. The same concentration of normal rabbit IgG (0.05 µg/mL, product number 2729; Cell Signaling Technology, Danvers, MA) was used as a negative control.

### Double-labeled immunofluorescence staining

Double-labeled immunofluorescence staining was performed as previously described.^20–22^ Briefly, deparaffinized and rehydrated sections of rat kidneys were unmasked by autoclaving for 5 minutes at 121°C in 1.0 mmol/L EDTA buffer (pH 9.0). After incubating with 5% BSA/PBS for 30 minutes at room temperature, the sections were reacted overnight at 4°C in a moist chamber with mixtures of primary antibodies against SGLT2 (Table 1) and proximal tubular markers (Supplementary Table S2). The following day, the slides were treated for 30 minutes at room temperature with fluorophore-conjugated secondary antibodies (Alexa Fluor 488 Goat anti-mouse IgG, product number A11029, and Alexa Fluor 555 Goat anti-rabbit IgG, product number A21430, both diluted in 1:1000; Invitrogen, Carlsbad, CA) in a light-shielded moist chamber. Hoechst 33342 (5.0 µg/mL, product number H3570; Invitrogen) was used for nuclear counterstaining. After autofluorescence was quenched using 0.1% Sudan black B (product number 199664; Sigma-Aldrich) in 70% ethanol,^23^ the sections were mounted with ProLong Gold Antifade mounting medium (product number P36930; Invitrogen) and sealed with a micro cover glass (NEO micro cover glass, 0.13-0.17 mm thickness; Matsunami Glass Industry). Immunofluorescence images were acquired by an Olympus SpinSR10 spinning disk confocal super resolution microscope (Evident, Tokyo, Japan) using an oil-immersion 60x objective (Olympus PlanApo N 60x/1.42 Oil Microscope Objective, Evident).

The tyramide signal amplification (TSA)-based multiplex immunofluorescence method was employed for double-labeling immunofluorescence staining using primary antibodies derived from the same host species, as previously described.^21,24^ Briefly, tissue sections were deparaffinized, retrieved antigens, and reacted overnight with the first rabbit primary antibody. Following incubation with a horseradish peroxidase (HRP)-conjugated secondary antibody (Histofine simple stain MAX-PO), covalent deposition of fluorophores around the recognition point of the first primary antibody was developed with the catalytic activity of horseradish peroxidase and iFluore 555 Styramide reagent (product number 45030; AAT Bioquest, Sunnyvale, CA). The binding between antibodies and target antigens was then stripped by autoclaving the sections for 5 minutes at 121°C in 10 mmol/L citrate buffer. Subsequently, the sections were incubated with the second rabbit primary antibody, followed by a fluorophore-conjugated secondary antibody (Alexa Fluor 488 anti-rabbit IgG, product number A11034, 1:1000), and counterstained with Hoechst 33342, as described above.

### Immunoelectron detection of DAB staining by low-vacuum scanning electron microscopy (LV-SEM)

Immunohistochemical DAB deposits were labeled with nanogold particles using an *in situ* nanogold labeling technique for scanning electron microscopy observation.^25^ Briefly, rat kidney sections that had undergone DAB staining for target proteins (SGLT2, LRP2, NHE3, and PDZK1IP1), as detailed in the preceding section, were treated with 0.01% tetrachloroauric acid (HAuCl_4_·4H_2_O, product number 077-00931; FUJIFILM Wako Pure Corporation) for 10 minutes at room temperature to promote nanogold nucleation at sites of DAB deposits. Following three rinses with distilled water, the slides were placed in a moist chamber at 37°C for 24 hours to develop the nucleated nanogold particles. The slides were then air-dried using a blower, and images were acquired using low-vacuum scanning electron microscopy (LV-SEM, Miniscope TM4000; Hitachi High-Technologies, Tokyo, Japan) under the following conditions: acceleration voltage of 15 kV and chamber pressure 30 Pa.

### Protein extraction

Frozen tissue samples were ground into a fine powder using a mortar and pestle pre-chilled with liquid nitrogen. The powdered tissue was divided into three portions and lysed in one of the following buffers: Cell Lysis Buffer (1x solution diluted with distilled water, product number 9803; Cell Signaling Technology), radioimmunoprecipitation assay (RIPA) buffer (1x solution diluted with distilled water, product number 9806; Cell Signaling Technology), or sodium dodecyl sulfate (SDS) lysis buffer (1% SDS, 1.0 mM EDTA, 10 mM Tris [pH 8.0]). Both Cell Lysis Buffer and RIPA buffer were supplemented with 1.0 mmol/L phenylmethylsulfonyl fluoride (product number 36978; Thermo Fisher Scientific, Waltham, MA) and a protease inhibitor cocktail (product number 11836153001; Roche, Basel, Switzerland). Lysates were sonicated for 30 seconds and centrifuged at 13,000 rpm for 5 minutes at 4°C. The supernatants were collected as total protein extracts. Protein concentrations were determined using the Pierce BCA Protein Assay Kit (product number 23225; Thermo Fisher Scientific).

### Western blotting

Twenty micrograms of total protein were mixed with Laemmli sample buffer (product number 1610747; Bio-Rad Laboratories, Hercules, CA) containing 5% (710 mM) 2-mercaptoethanol. To prevent aggregation of membrane proteins, 100 mM L-Arginine (A0526, Tokyo Chemical Industry, Tokyo, Japan), an inter-protein aggregation suppressor,^26^ was added prior to heat denaturation as appropriate. Samples were heat-denatured at 95°C for 5 minutes on a heat block incubator (WSC-2620 PowerBLOCK, product number 4002620; ATTO, Tokyo, Japan) and immediately cooled on ice.

To analyze glycosylation patterns, kidney tissue lysates prepared in SDS lysis buffer (20 μg of total protein) were mixed with Laemmli sample buffer, 5% 2-mercaptoethanol, 100 mM L-Arginine, and Glycoprotein Denaturing Buffer (1x containing 40 mM DTT and 0.5% SDS; New England Biolabs, Ipswich, MA), and then denatured at 100°C for 10 minutes. The denatured protein mixtures were incubated at 37°C for 4 hours with PNGase F (product number P0704; New England Biolabs) to remove N-linked glycans, or with *O*-Glycosidase (product number P0733; New England Biolabs) to remove O-linked glycans.

Protein samples, along with protein ladder (Precision Plus Protein Standard; Bio-Rad Laboratories), were separated by electrophoresis on a 4-15% polyacrylamide gel (Mini-PROTEAN TGX Precast Gel, product number 4561086; Bio-Rad Laboratories) using Tris-glycine running buffer (1x solution diluted with distilled water, product number 1610732; Bio-Rad Laboratories) at 150 V until the dye front reached the bottom of the gel. Proteins were then transferred onto a polyvinylidene difluoride (PVDF) membrane (Trans-Blot Turbo Midi PVDF Transfer Packs, product number 1704157; Bio-Rad Laboratories) using the Trans-Blot Turbo Transfer System (product number 1704150; Bio-Rad Laboratories) at 2.5 A and 25 V for 7 minutes. Membranes were blocked with PVDF Blocking Reagent for Can Get Signal (product number NYPBR01; TOYOBO, Osaka, Japan) for 30 minutes at room temperature, followed by overnight incubation at 4°C with primary antibodies against SGLT2 (1:1000; Table 1) or glyceraldehyde-3-phosphate dehydrogenase (GAPDH, 1:5000; Supplementary Table 2) diluted in Can Get Signal Solution 1 (product number NKB-201; TOYOBO) on a seesaw shaker (∼20 rpm, Wave-SI; TAITEC, Saitama, Japan). After three washes with Tris-buffered saline (TBS, 1x solution diluted with distilled water, product number 1706435; Bio-Rad Laboratories) containing 0.08% tween 20 (product number P1379; Sigma-Aldrich), membranes were incubated for 30 minutes at room temperature on a seesaw shaker (∼20 rpm) with HRP-conjugated secondary antibodies (1:5000, anti-mouse IgG, product number 7064, or anti-rabbit IgG, product number 7076; Cell Signaling Technology) diluted in Can Get Signal Solution 2 (product number NKB-301; TOYOBO). Protein bands were visualized using an enhanced chemiluminescence system (Clarity Max Western ECL Substrate, product number 1705062; Bio-Rad Laboratories), and chemiluminescent signals were captured with a digital imaging system (WSE-6300H LuminoGraph III; ATTO, Tokyo, Japan). Membranes were not stripped and re-probed, except when detecting the housekeeping gene, GAPDH, as a loading control.

## RESULTS

### Staining pattern of anti-SGLT2 antibodies

*Sglt1*- or *Sglt2*-deficient mice and rats were generated and exhibited glucosuria (Supplementary Figure S1-S4). Among the eight anti-SGLT2 antibodies tested, ab306558 and HPA041603 showed specific immunostaining in kidney sections from wild-type mice and rats when heat-induced antigen retrieval was performed using EDTA buffer (Figure 1 and Supplementary Figure S5-S12). Retrieval using citrate buffer resulted in weaker staining (Supplementary Figure S5-S12). In contrast, these two antibodies produced no SGLT2 immunostaining in cardiac sections (Supplementary Figure S13-S20). The remaining antibodies exhibited high background or non-specific staining in kidney sections, including vesicular-like deposits and glomerular staining (Supplementary Figure S21 and Supplementary Table S3).

**Figure 1.**
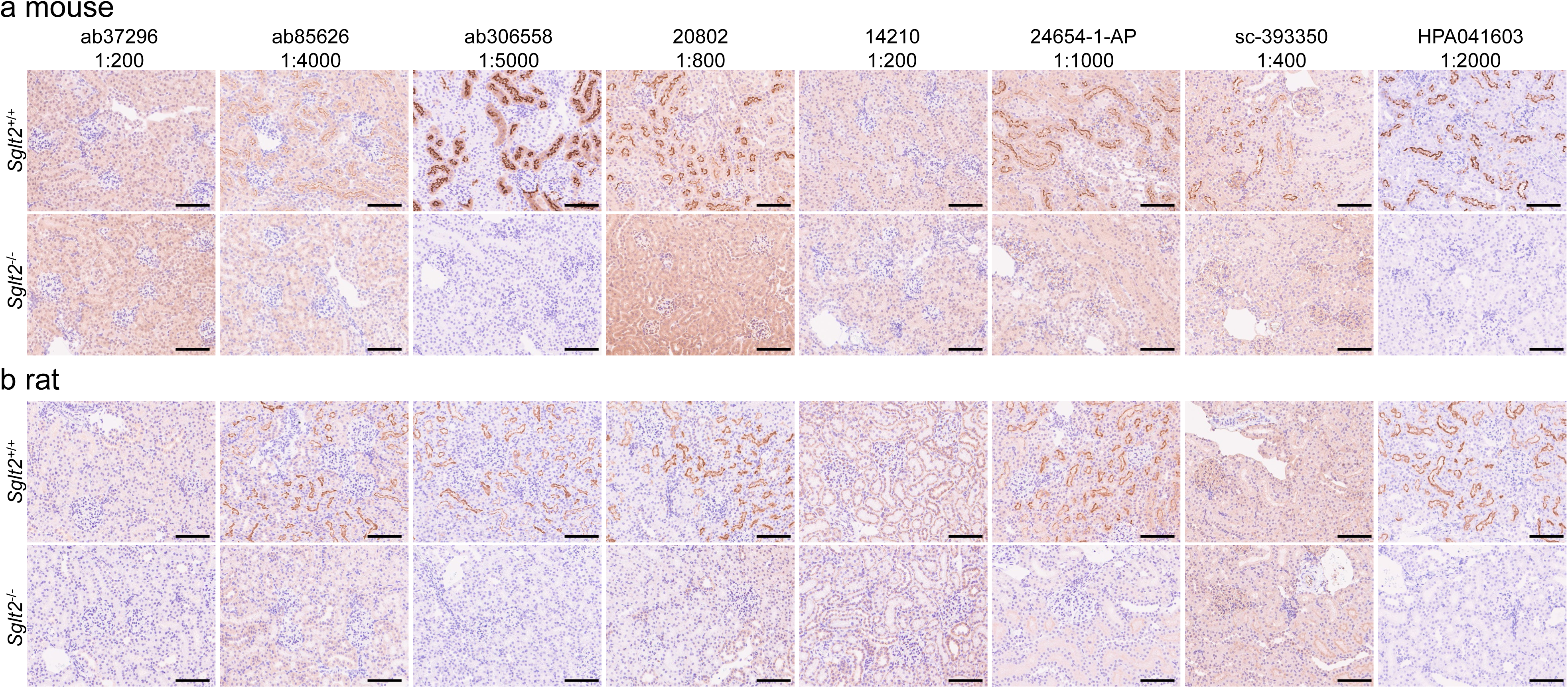
Staining patterns of eight anti-SGLT2 antibodies in wild-type and *Sglt2*-deficient rodent kidneys. (a, b) Serial kidney sections from wild-type (*Sglt2*^+/+^) and *Sglt2*-deficient (*Sglt2*^-/-^) mice (a) and rats (b) were immunostained with eight different anti-SGLT2 antibodies, each tested at five concentrations by sequential twofold dilutions (Supplementary Figure S5-S12). Representative images are shown for each antibody at the dilution with optimal signal-to-noise contrast. Among the antibodies, ab306558 and HPA041603 demonstrated SGLT2-specific staining in wild-type kidneys without background or non-specific staining in *Sglt2*-deficient sections. Nuclei were counterstained with hematoxylin. Scale bar = 100 μm.

The specificity of HPA041603 was confirmed by preincubation with its antigen peptide, which abolished immunostaining in human renal biopsy and rodent kidney sections (Figure 2a). No staining was observed with normal rabbit IgG (Figure 2a). In human renal cell carcinoma samples, SGLT2 immunoreactivity was present in the proximal tubules of non-tumor regions but absent in tumor areas (Figure 2b). In contrast, ab306558 showed inconsistent staining in human kidneys, with non-specific or absent signals (Supplementary Figure S22). Its specificity could not be assessed by the absorption test because the antigen sequence was undisclosed (Table 1). Based on these findings, HPA041603 was selected as the principal antibody for subsequent experiments, with ab306558 used for replication in rodents.

**Figure 2.**
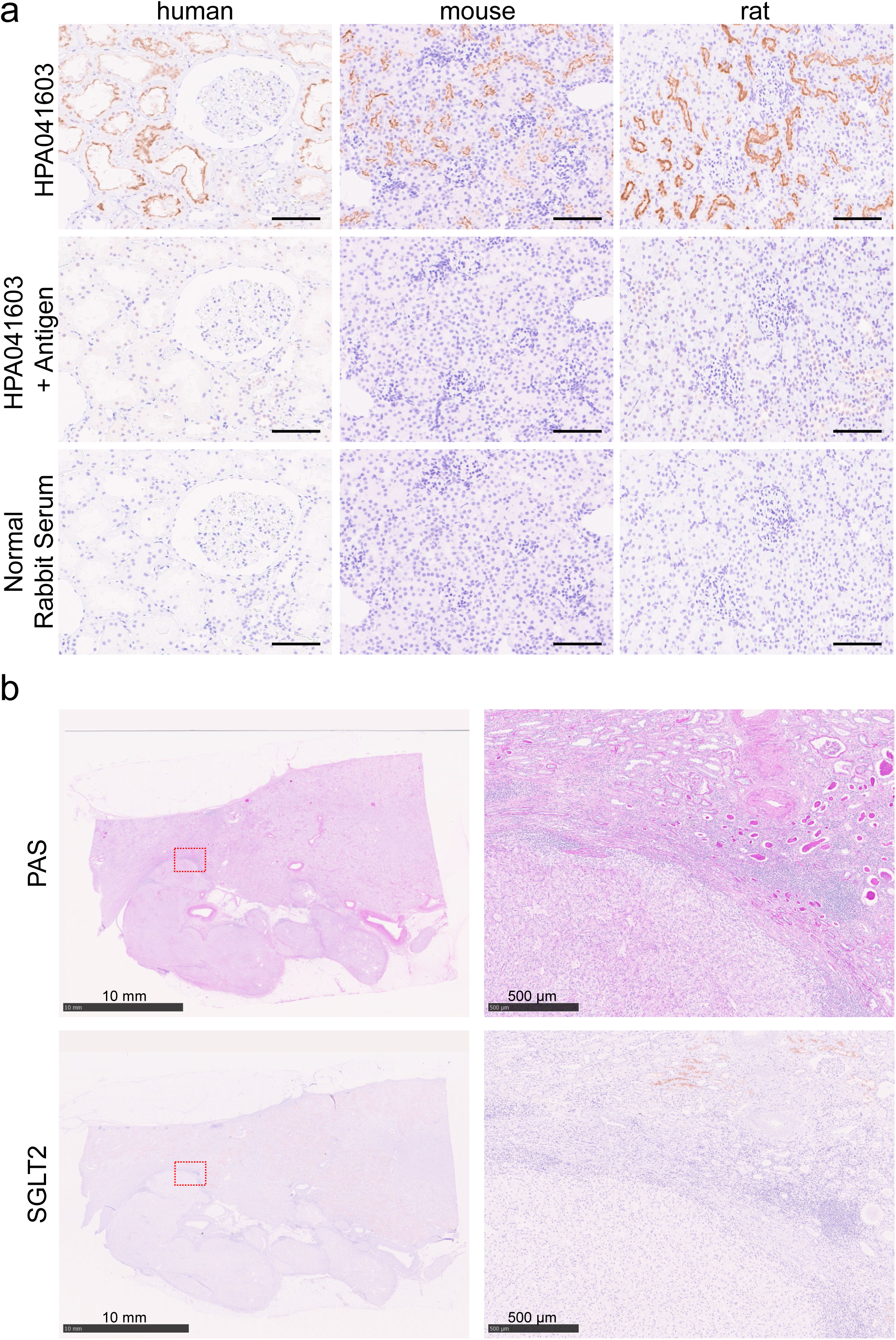
Absorption test and SGLT2 expression in human samples. (a) Absorption test for HPA041603 in human, mouse, and rat kidney sections. Specific SGLT2 immunoreactivity in renal proximal tubules was abolished after pre-absorption with the antigen peptide. No staining was observed with normal rabbit serum control. Nuclei were counterstained with hematoxylin. Scale bar = 100 μm. (b) Representative images of periodic acid-Schiff (PAS) staining and SGLT2 (HPA041603) immunostaining in kidney tissue from a patient with renal cell carcinoma (patient no. 4 in Supplementary Table S1). The boxed regions in the left panels are magnified in the right panels. SGLT2 immunoreactivity was detected in proximal tubules of non-tumor regions but was absent in carcinoma cells. Scale bars = 10 mm (left panels) and 500 μm (right panels).

### Localization of SGLT2 in rat proximal tubules

Immunostaining for PDZ domain containing 1 interacting protein 1 (PDZK1IP1, also known as membrane-associated protein 17 [MAP17]), an SGLT2 co-factor,^27^ was reduced in the proximal tubules of *Sglt2*-deficient rats compared with wild-type rats, while remaining detectable in infiltrating leukocytes (Supplementary Figure S23). The localization of low-density lipoprotein receptor-related protein 2 (LRP2, also known as megalin) and sodium/hydrogen exchanger 3 (NHE3) was unchanged between *Sglt2* genotypes (Supplementary Figure S23).

Low-vacuum scanning electron microscopy (LV-SEM) revealed that nanogold labeling for SGLT2 (HPA041603) and PDZK1IP1 was predominantly localized to microvilli at the apical membrane of proximal tubular cells, whereas LRP2 and NHE3 were distributed at the microvillar base (Figure 3a). Double-immunofluorescence staining confirmed colocalization of the HPA041603 signal with villin, a microvilli marker, but not with LRP2 or NHE3 (Figure 3b). PDZK1IP1 partially overlapped with the HPA041603 signal. Additionally, the HPA041603 signal did not overlap with acetylated alpha-tubulin, a primary cilia marker. These staining patterns were reproduced using ab306558 (Supplementary Figure S24).

**Figure 3.**
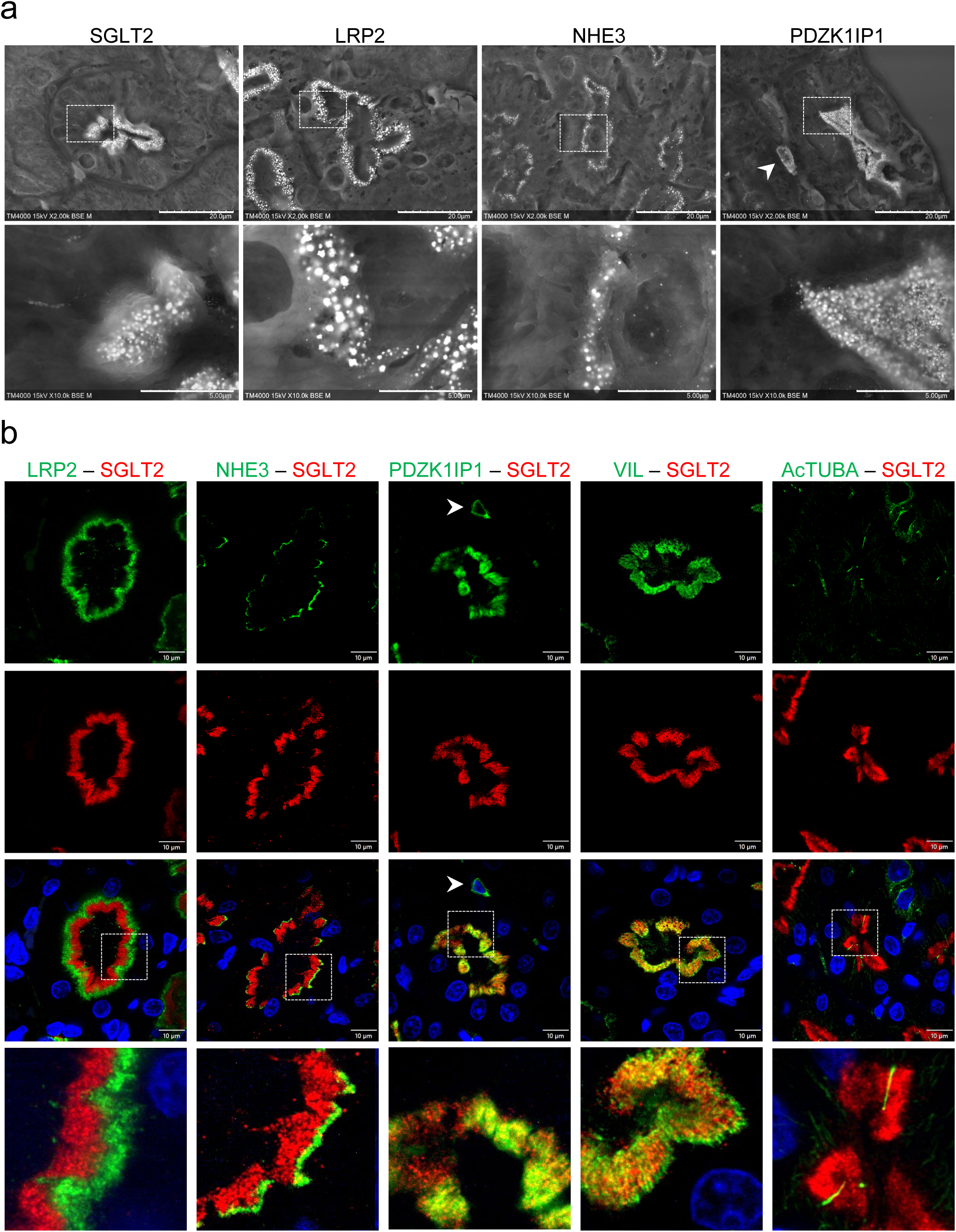
Subcellular localization of SGLT2 in rat kidney. (a) Representative low vacuum scanning electron microscopy (LV-SEM) images of SGLT2 (HPA041603), LRP2 (megalin), NHE3, and PDZK1IP1 (MAP17) in proximal tubular cells of wild-type rat kidneys. Arrowhead indicates PDZK1IP1-positive infiltrating blood cell. Top panels: scale bar = 20 μm (magnification ×2,000). Bottom panels: magnified views of boxed regions in the top panels. Scale bar = 5 μm (magnification ×10,000). (b) Representative double-immunofluorescence images of SGLT2 (HPA041603, red) with proximal tubular markers (green) in wild-type rat kidneys. SGLT2 shows partial overlap with PDZK1IP1, an SGLT2 co-factor, and colocalization with villin (VIL), a microvilli marker. LRP2 and NHE3 localize at the base of the microvilli and do not colocalize with SGLT2. Acetylated alpha-tubulin (AcTUBA), a primary cilia marker, also does not colocalize with SGLT2. Nuclei were counterstained with Hoechst 33342 (blue). Boxed regions in merged images are shown at higher magnification in the bottom panels. Arrowhead indicates PDZK1IP1-positive infiltrating blood cells. Scale bar = 10 μm.

### Western blotting for SGLT2 protein detection

When kidney lysates were prepared using Cell Lysis Buffer or radioimmunoprecipitation assay (RIPA) buffer, ab306558 and HPA041603 detected broad immunoreactive signals ranging from 50 to 300 kDa in wild-type and *Sglt1*-deficient samples, which were absent in *Sglt2*-deficient samples (Supplementary Figure S25). In contrast, SDS lysis buffer produced a single ∼55 kDa band with a slight smear. Supplementation with 100 mM L-arginine, a protein aggregation suppressor,^26^ partially reduced the smear without altering band migration (Supplementary Figure S25).

Based on these results, subsequent evaluation of the eight antibodies was performed using SDS lysis buffer supplemented with 100 mM L-arginine. A distinct band of ∼55 kDa was observed with ab306558, 20802, 24654-1-AP, and HPA041603 (Figure 4). Some antibodies showed non-specific bands, including signals near the predicted molecular mass of SGLT2 (∼73 kDa based on amino acid sequence). The ab85626 antibody detected a weak 55 kDa band even in *Sglt2*-deficient samples, suggesting cross-reactivity.

**Figure 4.**
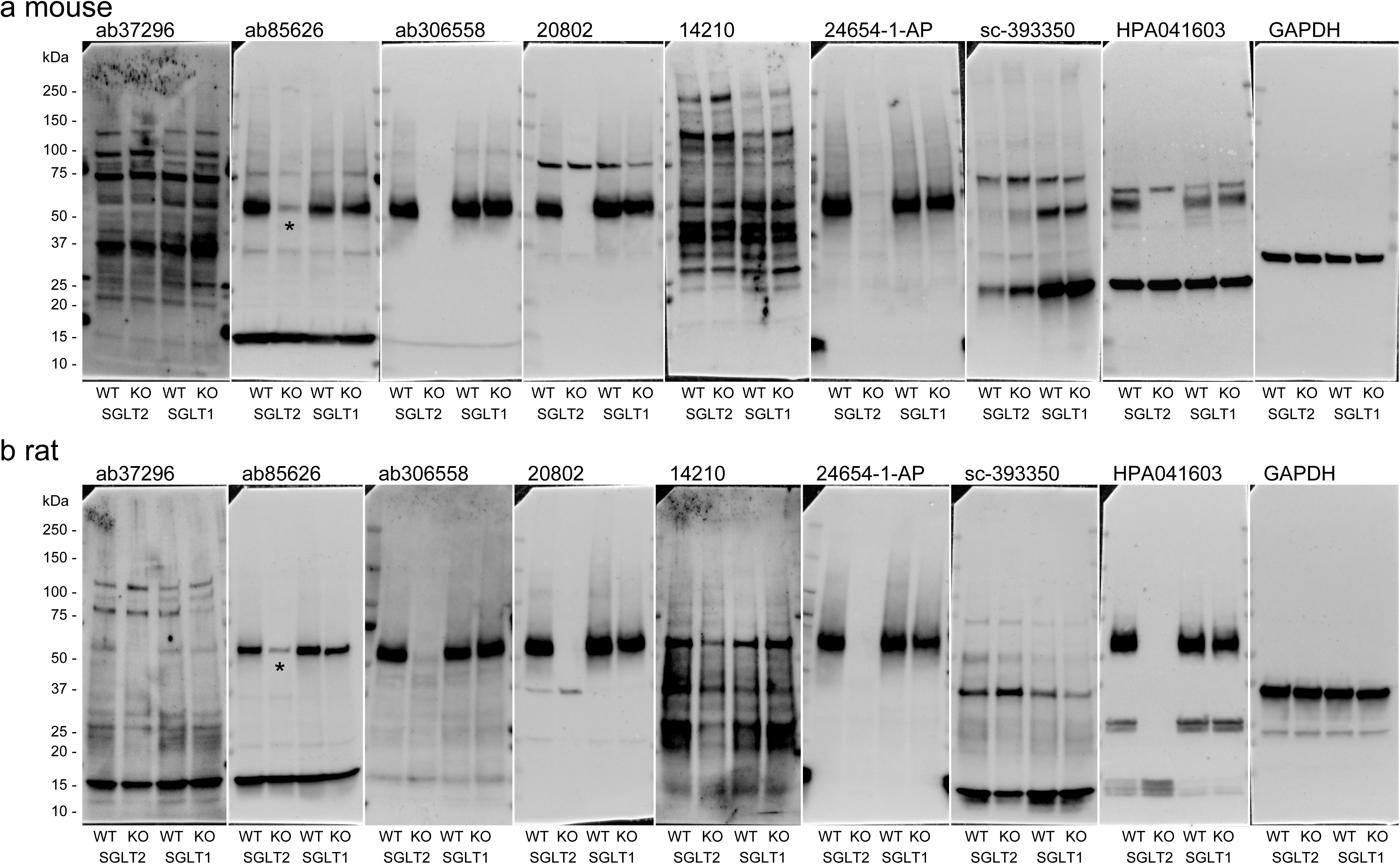
Western blotting of eight anti-SGLT2 antibodies in wild-type and *Sglt2*-deficient rodent kidneys. (a, b) Representative full Western blot images of whole kidney lysate from *Sglt2*-deficient mouse, *Sglt1*-deficient mouse, and their wild-type littermates (a), and kidney cortex lysates from *Sglt2*-deficient rats, *Sglt1*-deficient rats, and their wild-type littermates (b). SGLT2-specific bands were detected at approximately 55 kDa in wild-type and *Sglt1*-deficient samples. Some antibodies showed non-specific bands, including signals near 73 kDa, corresponding to the calculated molecular mass of SGLT2. Asterisks (*) indicate a weak 55 kDa band detected by ab85626 in *Sglt2*-deficient lysates, suggesting cross-reactivity with other proteins. Glyceraldehyde 3-phosphate dehydrogenase (GAPDH) was used as a loading control. Molecular masses based on the protein ladder are indicated on the left side of the blots. WT, wild-type; KO, knockout.

The consensus motif for N-linked glycosylation reported in human SGLT1^28^ and SGLT2^27^ was conserved across rodents (Figure 5a), supporting the use of deglycosylation analysis for band confirmation. PNGase F treatment shifted the SGLT2 band from ∼55 kDa to ∼45 kDa in kidney lysates, whereas *O*-glycosidase treatment had no effect (Figure 5b, c).

**Figure 5.**
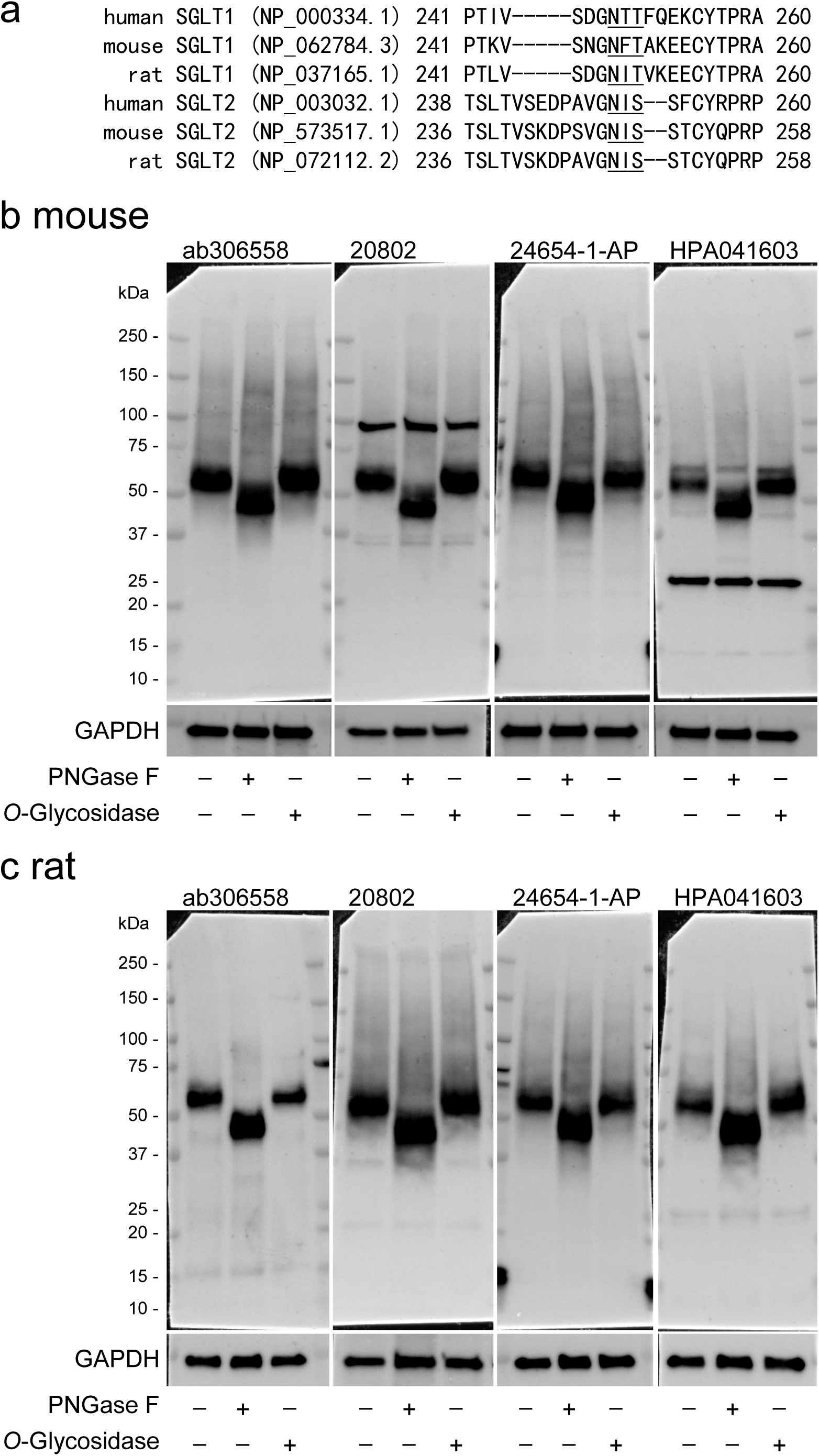
Effects of deglycosylation on SGLT2 band migration. (a) Multiple sequence alignment of SGLT1 and SGLT2 orthologs surrounding the predicted N-linked glycosylation motif, generated using the NCBI COBALT tool. The consensus sequence for N-linked glycosylation (N-X-T/S, where X is any amino acid except proline) is underlined. This glycosylation motif is conserved among human, mouse, and rat. (b, c) Representative full Western blot images of SGLT2 in wild-type kidney lysates from mouse (b) and rat (c) using selected anti-SGLT2 antibodies (ab306558, 20802, 24654-1-AP, and HPA041603). In untreated samples, a band at approximately 55 kDa, corresponding to glycosylated SGLT2, was observed. This band shifted to approximately 45 kDa following treatment with PNGase F, an enzyme that removes N-linked glycans. Treatment with *O*-Glycosidase, which removes O-linked glycans, did not affect SGLT2 migration. Glyceraldehyde 3-phosphate dehydrogenase (GAPDH) was used as a loading control. Molecular masses based on the protein ladder are indicated on the left side of the blots.

## DISCUSSION

The present study assessed the specificity of eight commercially available anti-SGLT2 antibodies. To the best of our knowledge, this is the first head-to-head comparison performed using genetically engineered *Sglt2*-deficient rodents as definitive negative controls. Among these antibodies, only a limited number provided reliable detection of SGLT2 protein with minimal background and without non-specific staining in immunohistochemistry. In Western blot analysis, the immunoreactive SGLT2 band appeared at approximately 55 kDa in rodent kidney lysate under optimized lysis buffer conditions, and this band shifted to about 45 kDa after enzymatic removal of N-linked glycans. These findings differed from the information provided in vendor datasheets (Table 1), reinforcing the need for rigorous and application-specific characterization of antibodies prior to protein expression studies.

Sample preparation procedures markedly influenced SGLT2 protein detection. EDTA-based antigen retrieval was superior to citrate buffer for unmasking SGLT2 epitopes in immunohistochemistry. In Western blotting, SDS lysis buffer minimized SGLT2 protein aggregation compared with Cell Lysis Buffer or RIPA buffer. These results highlight that protocol optimization is critical for accurate detection of the SGLT2 protein. However, the widespread use of both techniques has resulted in diverse methodologies, including unique laboratory adaptations^29,30^ and the incomplete reporting of procedural details as routine methods in the literature,^8^ thereby compromising reproducibility.

SGLT2 immunostaining was observed in the microvilli of proximal tubular epithelial cells in wild-type rats. This microvillar distribution likely maximizes the surface area of the transporter exposed to urinary glucose for efficient reabsorption. SGLT2 and PDZK1IP1 partially colocalized in wild-type rats, and PDZK1IP1 immunostaining in the microvilli was diminished in *Sglt2*-deficient rats. Interaction of PDZK1IP1 with SGLT2 has been reported to induce conformational changes that enable glucose and sodium transport.^27^ Because SGLT2 operates at approximately 50% of maximal capacity under physiological conditions,^31^ this partial overlap may represent a regulatory mechanism that fine-tunes transporter activity in response to physiological demand.

The observed bands (∼55 kDa glycosylated and ∼45 kDa after deglycosylation) were lower than the theoretical molecular mass of SGLT2 (∼73 kDa) in Western blot analysis. This phenomenon, known as “gel shifting”, commonly occurs in membrane proteins and is attributed to altered SDS/protein complex stoichiometry.^32^ SGLT1, a homolog of SGLT2 with a comparable calculated mass (∼73 kDa), also migrates at approximately 55 kDa.^33,34^ Furthermore, some antibodies detected non-specific bands near 73 kDa (Figure 4), emphasizing that careful discrimination between specific and non-specific bands is essential when interpreting SGLT2 presence in Western blot analyses. Enzymatic deglycosylation assays could be a beneficial tool to confirm the SGLT2 band.

Among the antibodies tested, ab85626 is one of the most frequently cited for SGLT2 detection across multiple species and applications in the CiteAb (https://www.citeab.com/). However, it exhibited a weak 55 kDa band even in *Sglt2*-deficient kidney lysates, suggesting cross-reactivity with other SLC5A family members or unrelated proteins. The linear epitopes of antibodies typically range from 4 to 12 amino acids.^35^ This length may allow specific target recognition, but it also raises the risk of cross-reactivity, particularly for polyclonal antibodies or antibodies against conserved protein families. Without appropriate knockout controls, such cross-reactivity could lead to misattribution of SGLT2 expression, especially in non-renal or malignant tissues where SGLT2 expression has been proposed.^5,36^ Indeed, immunostaining with HPA041603, a well-validated antibody in this study, showed no detectable SGLT2 in renal carcinoma cells.

In conclusion, this study demonstrates substantial variability in the specificity of commercially available anti-SGLT2 antibodies. Only a limited number are suitable for reliable detection of SGLT2 protein in rodent and human tissues under optimized procedures. Antibodies are indispensable tools in biomedical research, but improper use or insufficient validation can result in compromised data and irreproducible outcomes.^6–8^ Unlike therapeutic antibodies which are comprehensively characterized and regulated, antibodies used in the research field lack universal quality criteria.^37^ Rigorous antibody validation is essential to ensure reliable and reproducible findings and to avoid misinterpretation in basic and translational research involving membrane proteins such as SGLT2.

## Data Statement

All data generated or analyzed in this study were included in the main text and the Supplementary Material for this article. Any additional data that support the findings of this study are available from the corresponding author upon reasonable request.

## Acknowledgments

We would like to thank Dr. Makoto Matsuyama for his technical assistance with the *i*-GONAD procedure. We are also grateful to the Division of Diagnostic Pathology of Tohoku Medical and Pharmaceutical University Hospital for their support in handling human kidney samples. We acknowledge the Center for Laboratory Animal Science of Tohoku Medical and Pharmaceutical University for assistance with animal care and housing, and the Technical Services Department and the Histopathology Core Facility of Tohoku Medical and Pharmaceutical University for their support in histological sample preparation.

This study was supported in part by Grants-in-Aid for Scientific Research (20K08612, 23K07690, 24K10956, 25K11324, and 25K19425) from the Ministry of Education, Culture, Sports, Science, and Technology of Japan (MEXT), the Takeda Research Foundation, the Mochida Memorial Foundation for Medical and Pharmaceutical Research, the Kowa Life Science Foundation, and the Miyagi Kidney Foundation.

## Author Contributions

TH and MT conceived and designed the study. TH, HI, AE, CT, RI, and AK generated *Sglt2*- and *Sglt1*-deficient rodents using the *i*-GONAD method. HI, AE, TK, KY, RI, AK, IOY, MS, KM, TK, YN, and YW collected and processed human kidney biopsy, autopsy, and nephrectomy samples. TH, HI, AE, and TK performed immunohistochemistry. TH, TK, and KK carried out low-vacuum scanning electron microscopy and double-immunofluorescence imaging. TH, SS, CT, and RI conducted Western blotting and deglycosylation assays. TH drafted the manuscript. All authors contributed to data interpretation, critically revised the manuscript, and approved the final version.

