## Supplementary Material for "Reliable Detection of SGLT2 Protein through Knockout-based Antibody Characterization and Optimized Procedures"

### Supplementary Figure S1

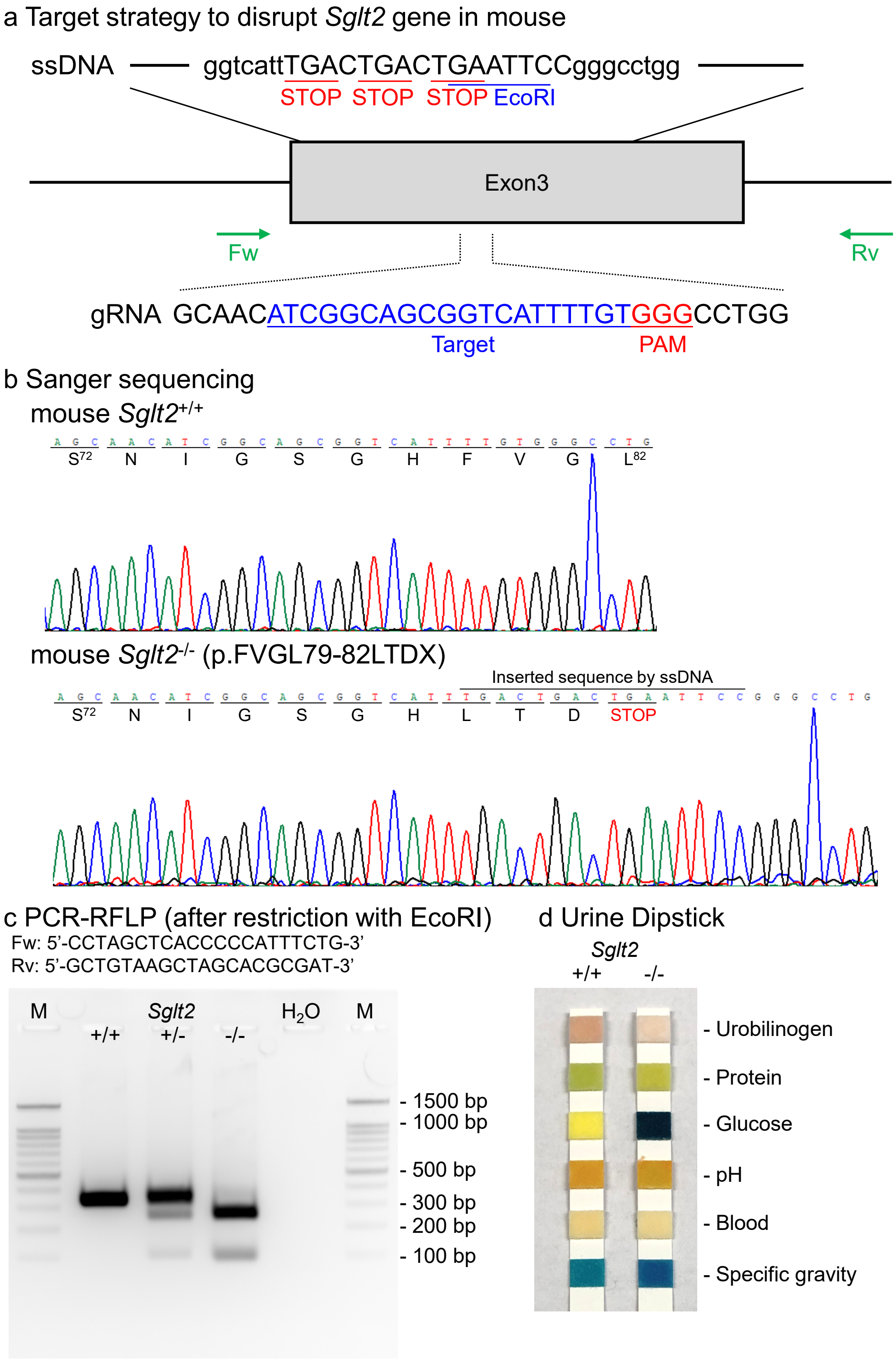


**Supplementary Figure S1. Generation of *Slc5a2*-deficient mice.**
(a) Schematic diagram of mouse *Slc5a2* gene locus illustrating the target sequence of single-strand oligodeoxynucleotide (ssODN), guide RNA (gRNA), and protospacer adjacent motif (PAM). Fw; forward primer, Rv; reverse primer for genotyping. (b) Representative Sanger sequencing chromatograms confirming genotypes of *Slc5a2* wild-type (*Slc5a2*$^{+/+}$) and knockout (*Slc5a2*$^{-/-}$) mice. (c) Representative gel image of polymerase chain reaction-restriction fragment length polymorphism (PCR-RFLP) genotyping following EcoRI digestion. The 305 base pairs (bp) fragment is present in *Slc5a2*$^{+/+}$ mice; 305 bp, 232 bp, and 85 bp fragments are found in *Slc5a2*$^{+/-}$ heterozygous mice; and 232 and 85 bp fragments are observed in *Slc5a2*$^{-/-}$ mice. M; molecular size marker. (d) Dipstick urinalysis showing glucosuria in *Slc5a2*$^{-/-}$ mice.

### Supplementary Figure S2

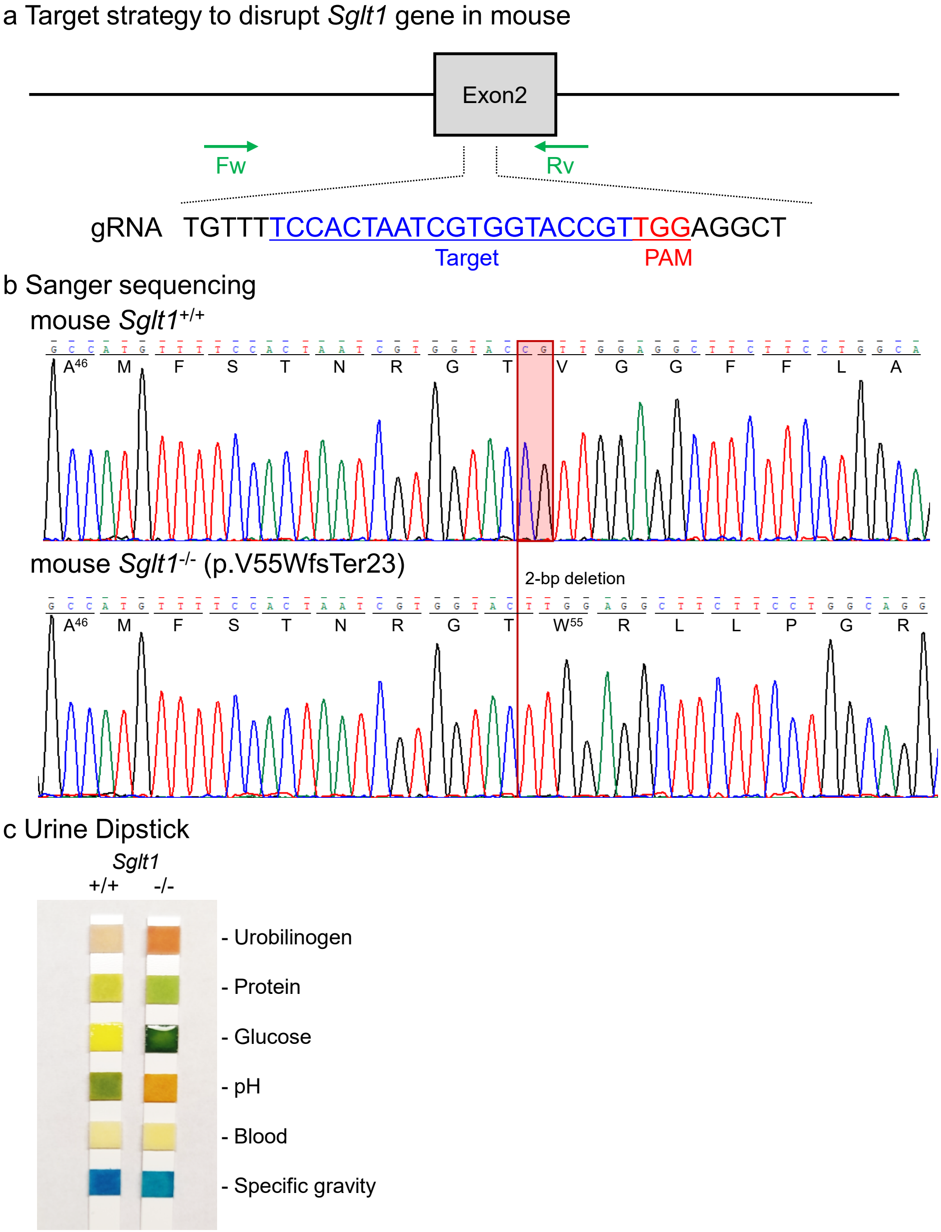


**Supplementary Figure S2. Generation of *Slc5a1*-deficient mice.**
(a) Schematic diagram of mouse *Slc5a1* gene locus illustrating the target sequence of guide RNA (gRNA), and protospacer adjacent motif (PAM). Fw; forward primer, Rv; reverse primer for Sanger sequencing. (b) Representative Sanger sequencing chromatograms confirming genotypes of *Slc5a1* wild-type (*Slc5a1*$^{+/+}$) and knockout (*Slc5a1*$^{-/-}$) mice. (c) Dipstick urinalysis showing glucosuria in *Slc5a1*$^{-/-}$ mice.

**Supplementary Figure S3**

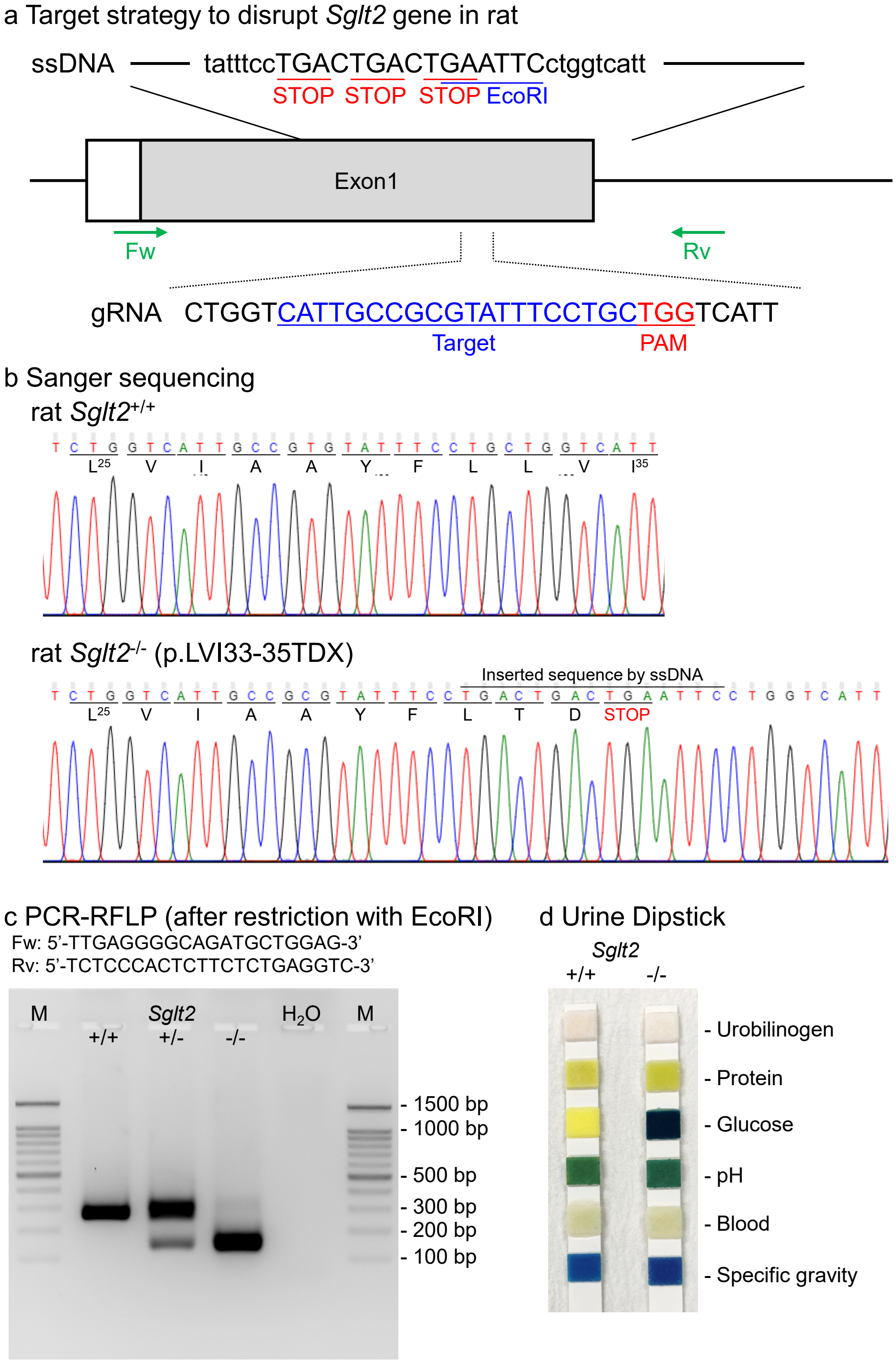


**Supplementary Figure S3. Generation of *Slc5a2*-deficient rats.**
(a) Schematic diagram of rat *Slc5a2* gene locus illustrating the target sequence of single-strand oligodeoxynucleotide (ssODN), guide RNA (gRNA), and protospacer adjacent motif (PAM). Fw; forward primer, Rv; reverse primer for genotyping. (b) Representative Sanger sequencing chromatograms confirming genotypes of *Slc5a2* wild-type (*Slc5a2*$^{+/+}$) and knockout (*Slc5a2*$^{-/-}$) rats. (c) Representative gel image of polymerase chain reaction-restriction fragment length polymorphism (PCR-RFLP) genotyping following EcoRI digestion. The 234 base pairs (bp) fragment is present in *Slc5a2*$^{+/+}$ rats; 234 bp, 129 bp, and 118 bp fragments are found in *Slc5a2*$^{+/-}$ heterozygous rats; and 129 and 118 bp fragments are observed in *Slc5a2*$^{-/-}$ rats. M; molecular size marker. (d) Dipstick urinalysis showing glucosuria in *Slc5a2*$^{-/-}$ rats.

### Supplementary Figure S4

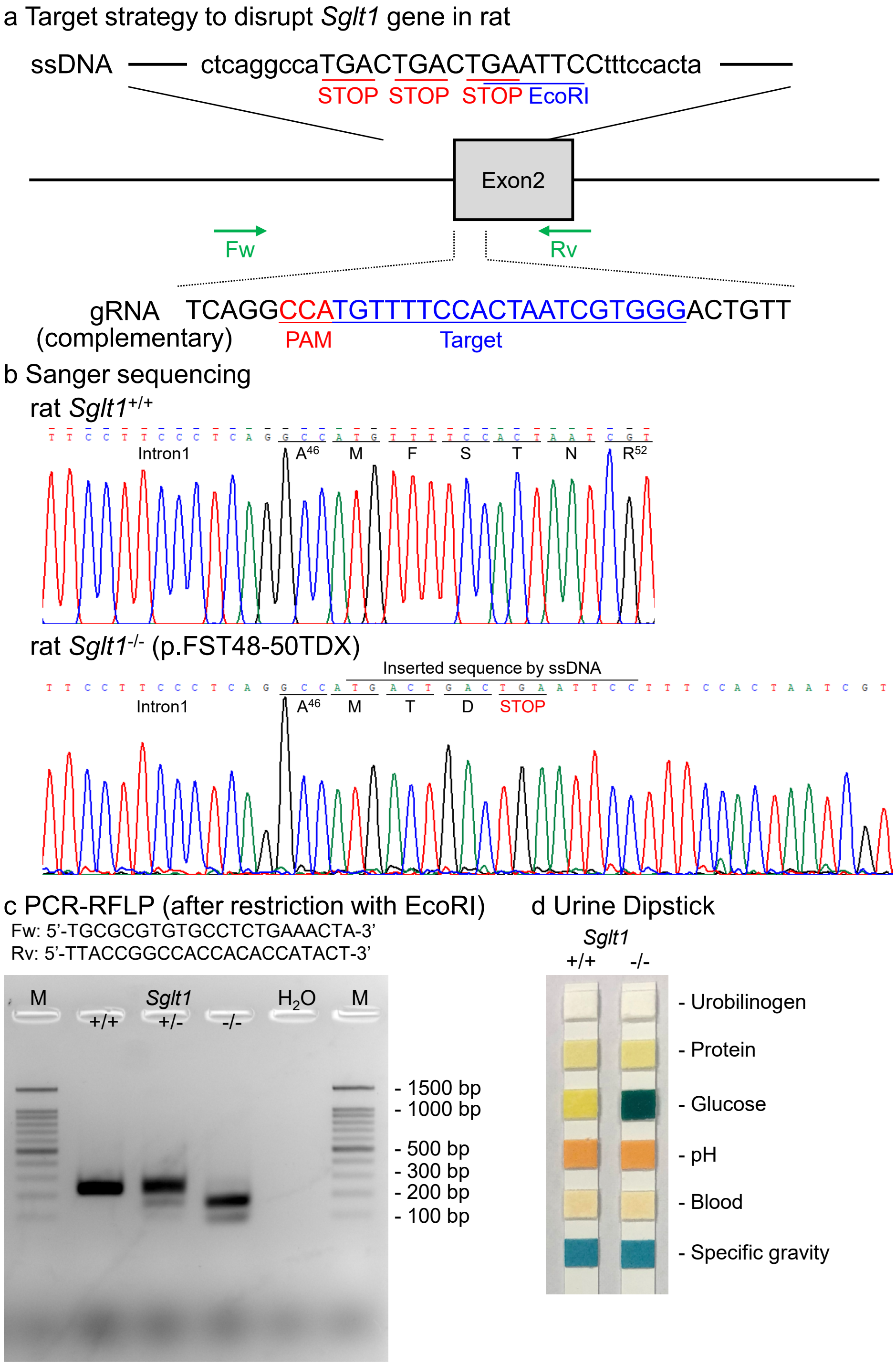


**Supplementary Figure S4. Generation of *Slc5a1*-deficient rats.**
(a) Schematic diagram of rat *Slc5a1* gene locus illustrating the target sequence of single-strand oligodeoxynucleotide (ssODN), guide RNA (gRNA), and protospacer adjacent motif (PAM). Fw; forward primer, Rv; reverse primer for genotyping. (b) Representative Sanger sequencing chromatograms confirming genotypes of *Slc5a1* wild-type ($Slc5a1^{+/+}$) and knockout ($Slc5a1^{-/-}$) rats. (c) Representative gel image of polymerase chain reaction-restriction fragment length polymorphism (PCR-RFLP) genotyping following EcoRI digestion. The 200 base pairs (bp) fragment is present in $Slc5a1^{+/+}$ rats; 200 bp, 140 bp, and 73 bp fragments are found in $Slc5a1^{+/-}$ heterozygous rats; and 140 and 73 bp fragments are observed in $Slc5a1^{-/-}$ rats. M; molecular size marker. (d) Dipstick urinalysis showing glucosuria in $Slc5a1^{-/-}$ rats.

### Supplementary Figure S5

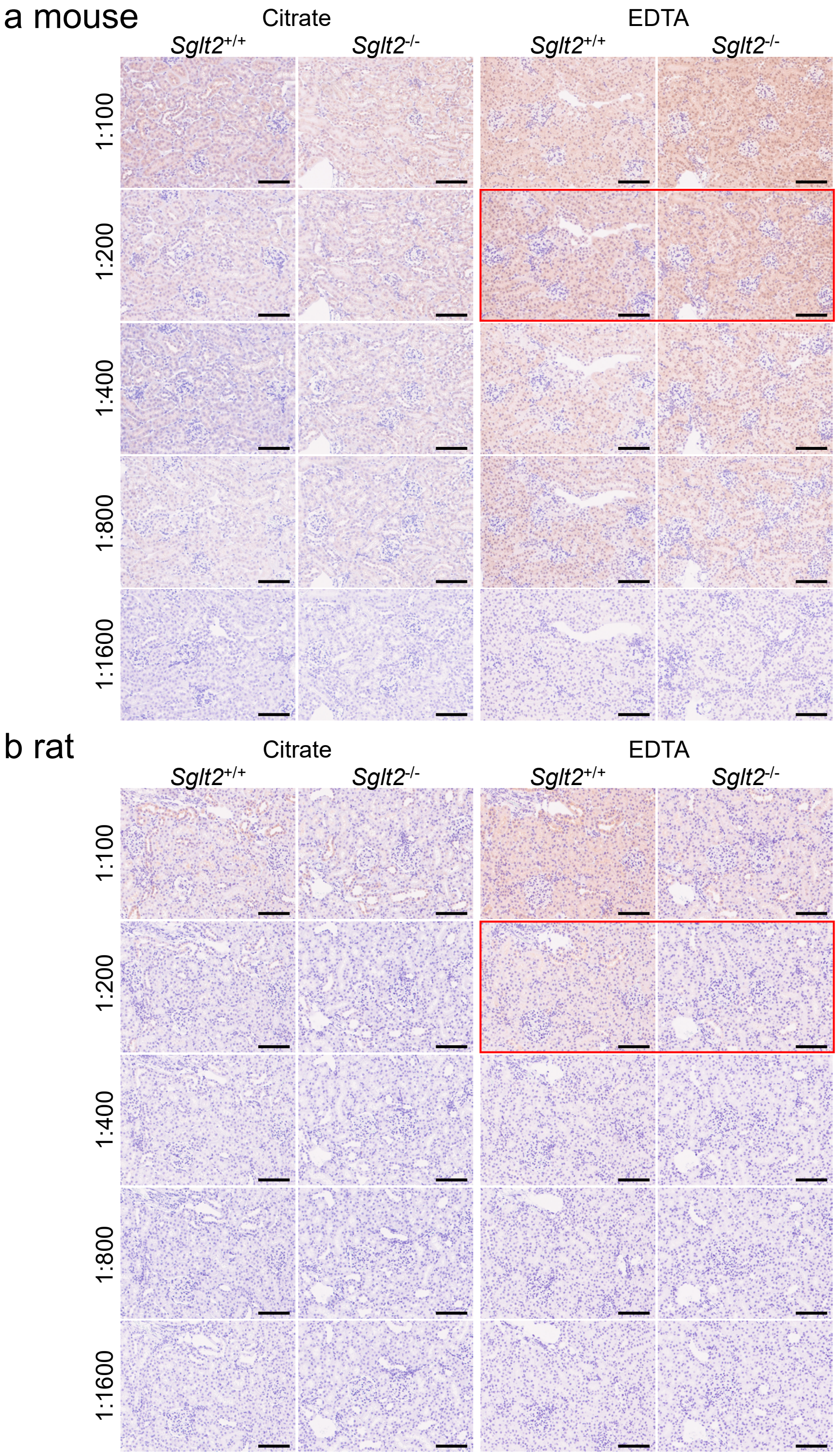


**Supplementary Figure S5. Optimization of anti-SGLT2 (ab37296) antibody concentration and antigen retrieval in rodent kidney sections.**
Chromogenic detection of anti-SGLT2 (ab37296) antibody immunostaining in the cortical regions of kidneys from wild-type (*Sglt2*$^{+/+}$) and *Sglt2*-deficient (*Sglt2*$^{-/-}$) mice (a) and rats (b). Five antibody dilutions (1:100 to 1:1600, twofold serial dilutions) and two antigen retrieval solutions (citrate buffer, pH 6.0 and EDTA buffer, pH 9.0) were tested. Nuclei were counterstained with hematoxylin. Images were captured using NanoZoomer-SQ slide scanner and exported to JPG file at x20 magnification using NDP.view2 software. The red box indicates the photo shown in Figure 1. Scale bar = 100 µm.

**Supplementary Figure S6**

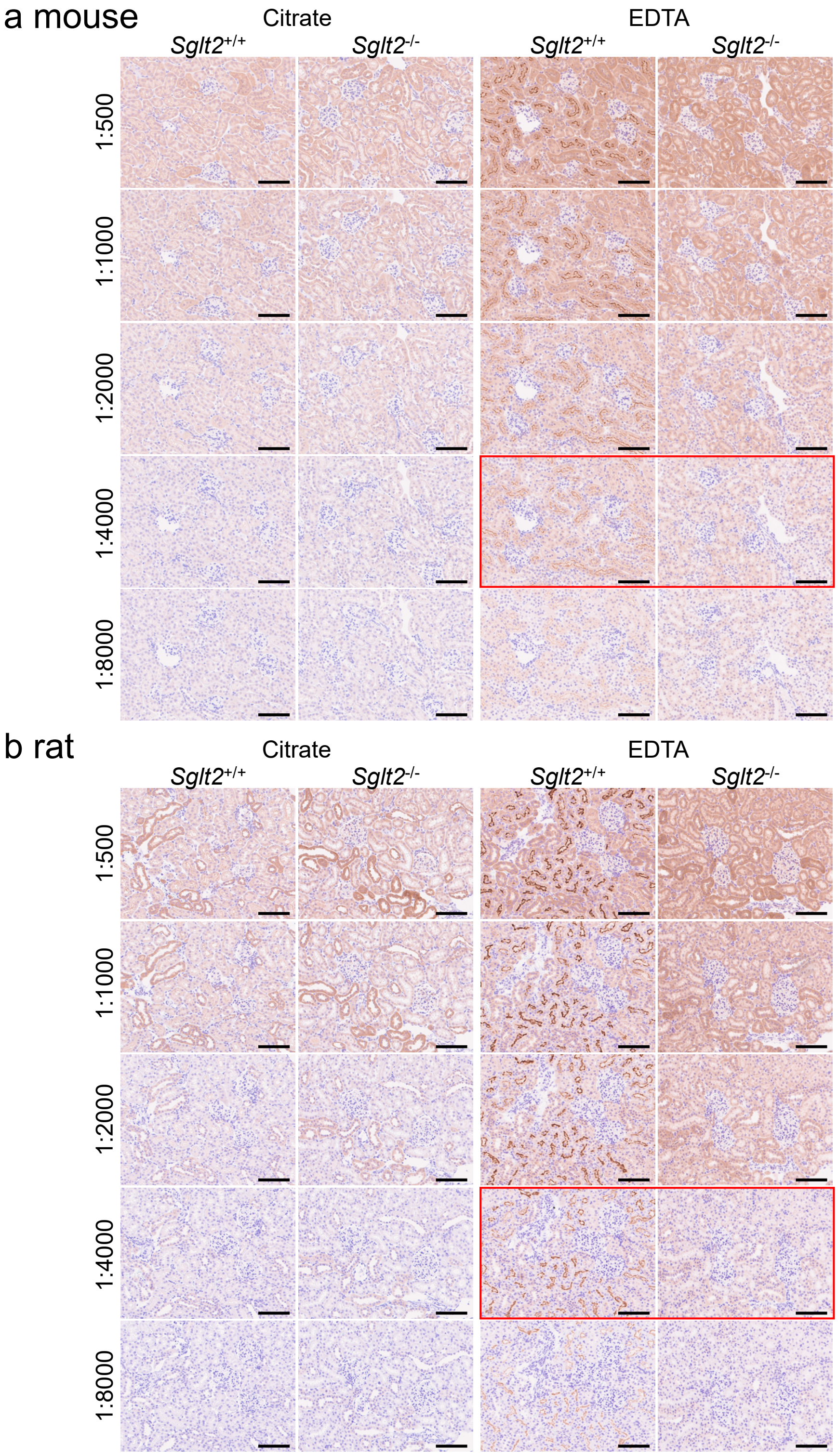


**Supplementary Figure S6. Optimization of anti-SGLT2 (ab85626) antibody concentration and antigen retrieval in rodent kidney sections.**
Chromogenic detection of anti-SGLT2 (ab85626) antibody immunostaining in the cortical regions of kidneys from wild-type (*Sglt2*$^{+/+}$) and *Sglt2*-deficient (*Sglt2*$^{-/-}$) mice (a) and rats (b). Five antibody dilutions (1:500 to 1:8000, twofold serial dilutions) and two antigen retrieval solutions (citrate buffer, pH 6.0 and EDTA buffer, pH 9.0) were tested. Nuclei were counterstained with hematoxylin. Images were captured using NanoZoomer-SQ slide scanner and exported to JPG file at x20 magnification using NDP.view2 software. The red box indicates the photo shown in Figure 1. Scale bar = 100 µm.

**Supplementary Figure S7**

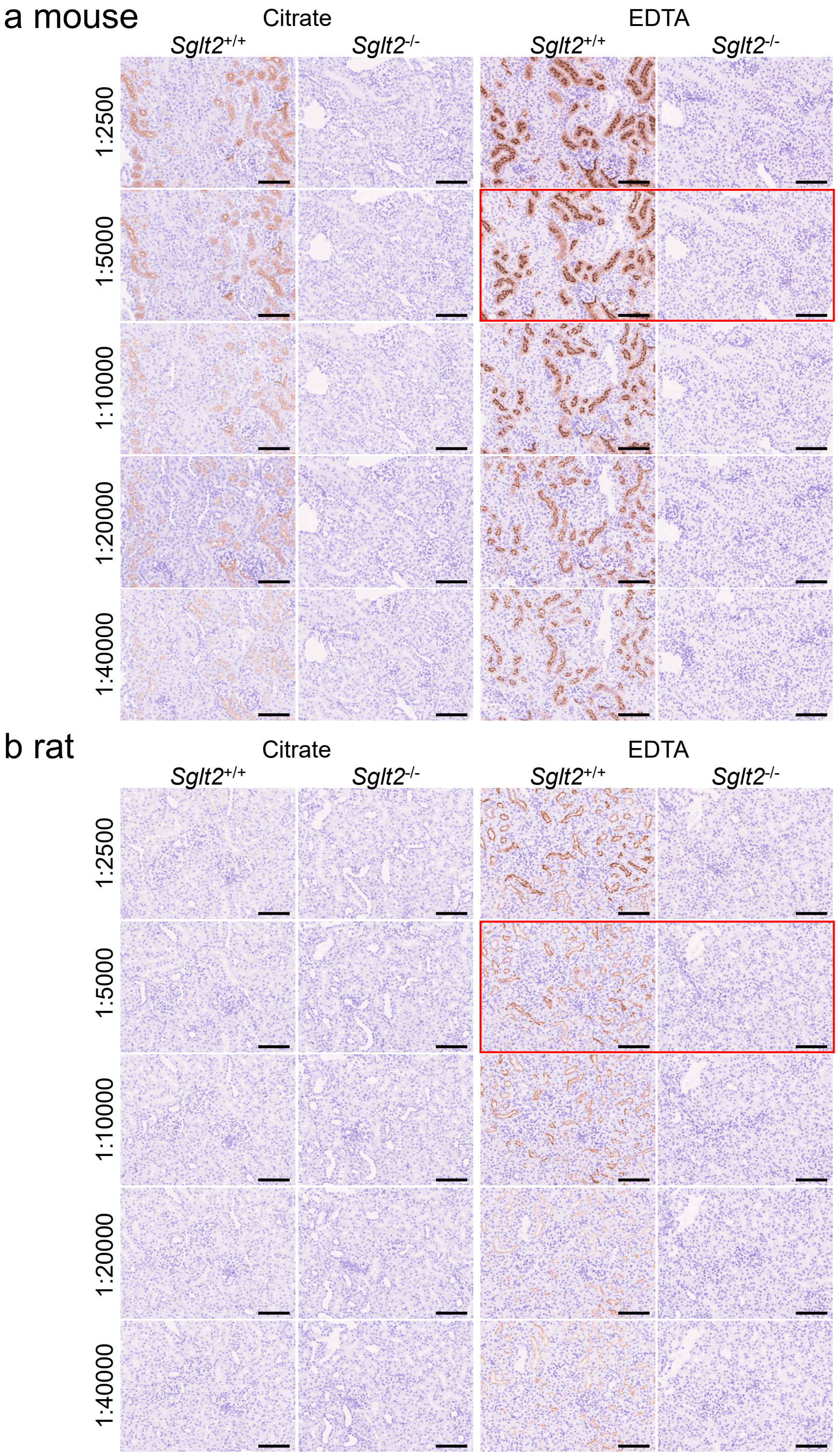


**Supplementary Figure S7. Optimization of anti-SGLT2 (ab306558) antibody concentration and antigen retrieval in rodent kidney sections.**
Chromogenic detection of anti-SGLT2 (ab306558) antibody immunostaining in the cortical regions of kidneys from wild-type (*Sglt2*$^{+/+}$) and *Sglt2*-deficient (*Sglt2*$^{-/-}$) mice (a) and rats (b). Five antibody dilutions (1:2500 to 1:4000, twofold serial dilutions) and two antigen retrieval solutions (citrate buffer, pH 6.0 and EDTA buffer, pH 9.0) were tested. Nuclei were counterstained with hematoxylin. Images were captured using NanoZoomer-SQ slide scanner and exported to JPG file at x20 magnification using NDP.view2 software. The red box indicates the photo shown in Figure 1. Scale bar = 100 µm.

**Supplementary Figure S8**

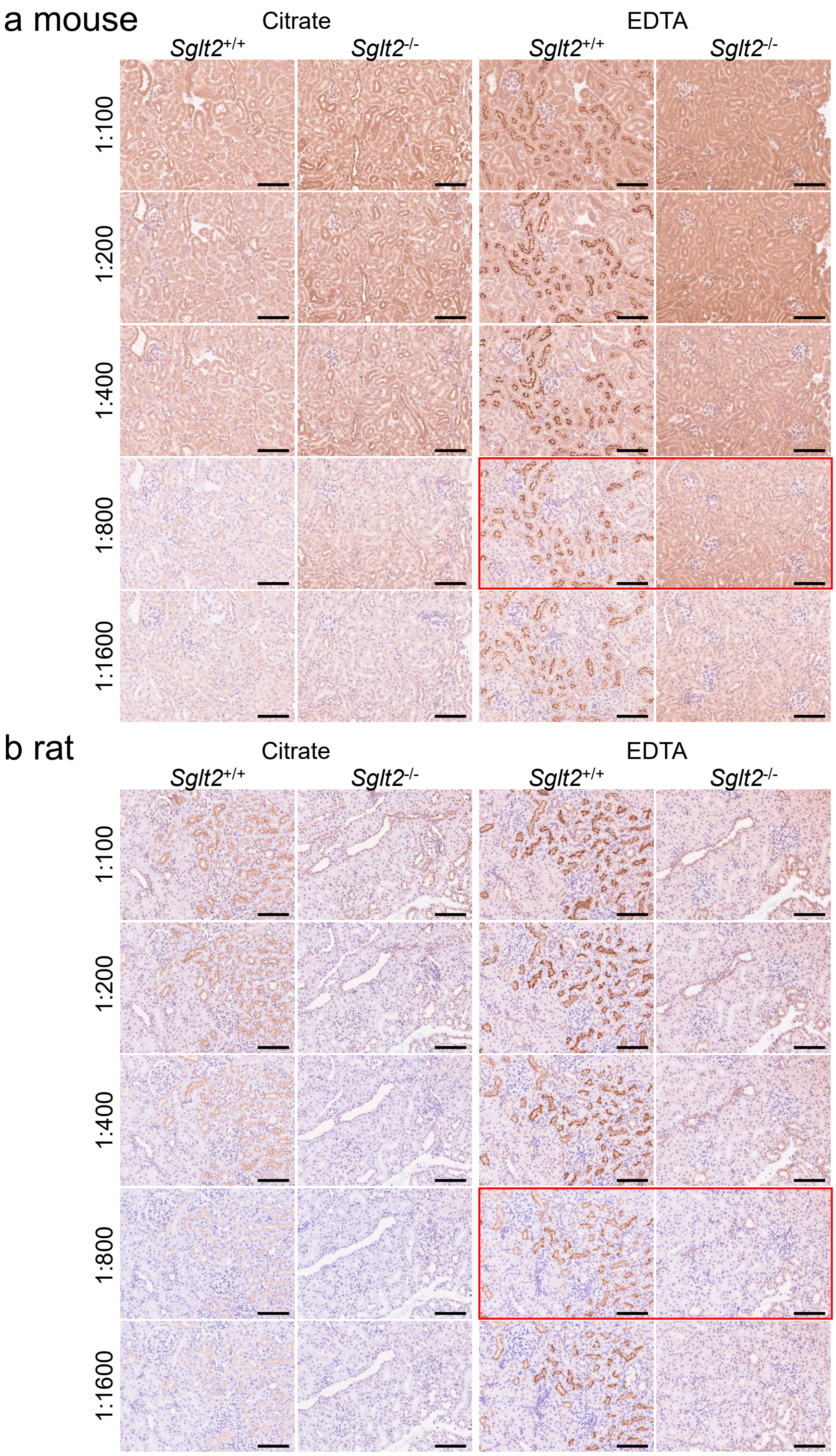


**Supplementary Figure S8. Optimization of anti-SGLT2 (20802) antibody concentration and antigen retrieval in rodent kidney sections.**
Chromogenic detection of anti-SGLT2 (20802) antibody immunostaining in the cortical regions of kidneys from wild-type (*Sglt2*$^{+/+}$) and *Sglt2*-deficient (*Sglt2*$^{-/-}$) mice (a) and rats (b). Five antibody dilutions (1:100 to 1:1600, twofold serial dilutions) and two antigen retrieval solutions (citrate buffer, pH 6.0 and EDTA buffer, pH 9.0) were tested. Nuclei were counterstained with hematoxylin. Images were captured using NanoZoomer-SQ slide scanner and exported to JPG file at x20 magnification using NDP.view2 software. The red box indicates the photo shown in Figure 1. Scale bar = 100 µm.

**Supplementary Figure S9**

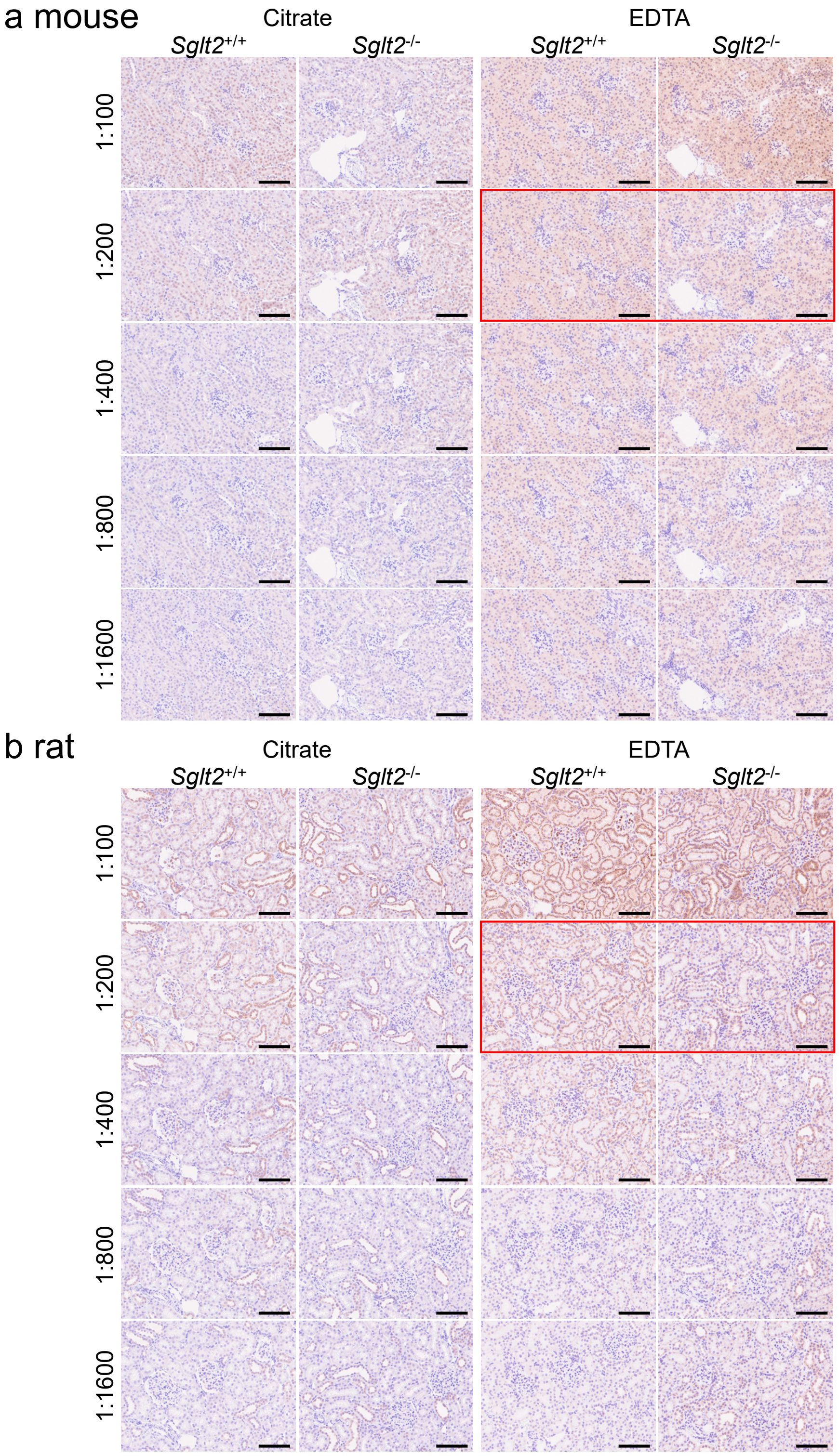


**Supplementary Figure S9. Optimization of anti-SGLT2 (14210) antibody concentration and antigen retrieval in rodent kidney sections.**
Chromogenic detection of anti-SGLT2 (14210) antibody immunostaining in the cortical regions of kidneys from wild-type (*Sglt2*$^{+/+}$) and *Sglt2*-deficient (*Sglt2*$^{-/-}$) mice (a) and rats (b). Five antibody dilutions (1:100 to 1:1600, twofold serial dilutions) and two antigen retrieval solutions (citrate buffer, pH 6.0 and EDTA buffer, pH 9.0) were tested. Nuclei were counterstained with hematoxylin. Images were captured using NanoZoomer-SQ slide scanner and exported to JPG file at x20 magnification using NDP.view2 software. The red box indicates the photo shown in Figure 1. Scale bar = 100 µm.

**Supplementary Figure S10**

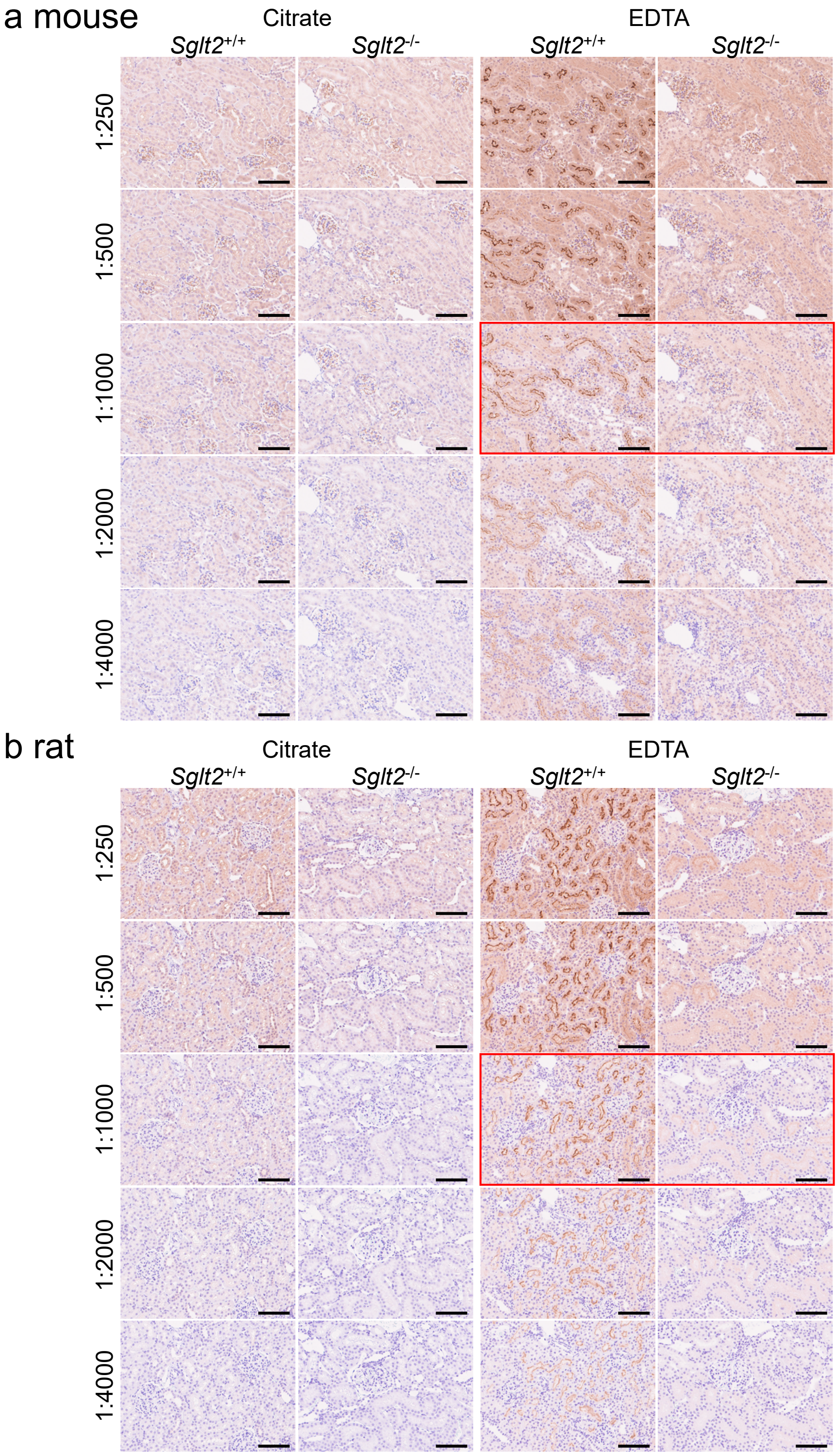


**Supplementary Figure S10. Optimization of anti-SGLT2 (24654-1-AP) antibody concentration and antigen retrieval in rodent kidney sections.**

Chromogenic detection of anti-SGLT2 (24654-1-AP) antibody immunostaining in the cortical regions of kidneys from wild-type (*Sglt2*$^{+/+}$) and *Sglt2*-deficient (*Sglt2*$^{-/-}$) mice (a) and rats (b). Five antibody dilutions (1:250 to 1:4000, twofold serial dilutions) and two antigen retrieval solutions (citrate buffer, pH 6.0 and EDTA buffer, pH 9.0) were tested. Nuclei were counterstained with hematoxylin. Images were captured using NanoZoomer-SQ slide scanner and exported to JPG file at x20 magnification using NDP.view2 software. The red box indicates the photo shown in Figure 1. Scale bar = 100 µm.

**Supplementary Figure S11**

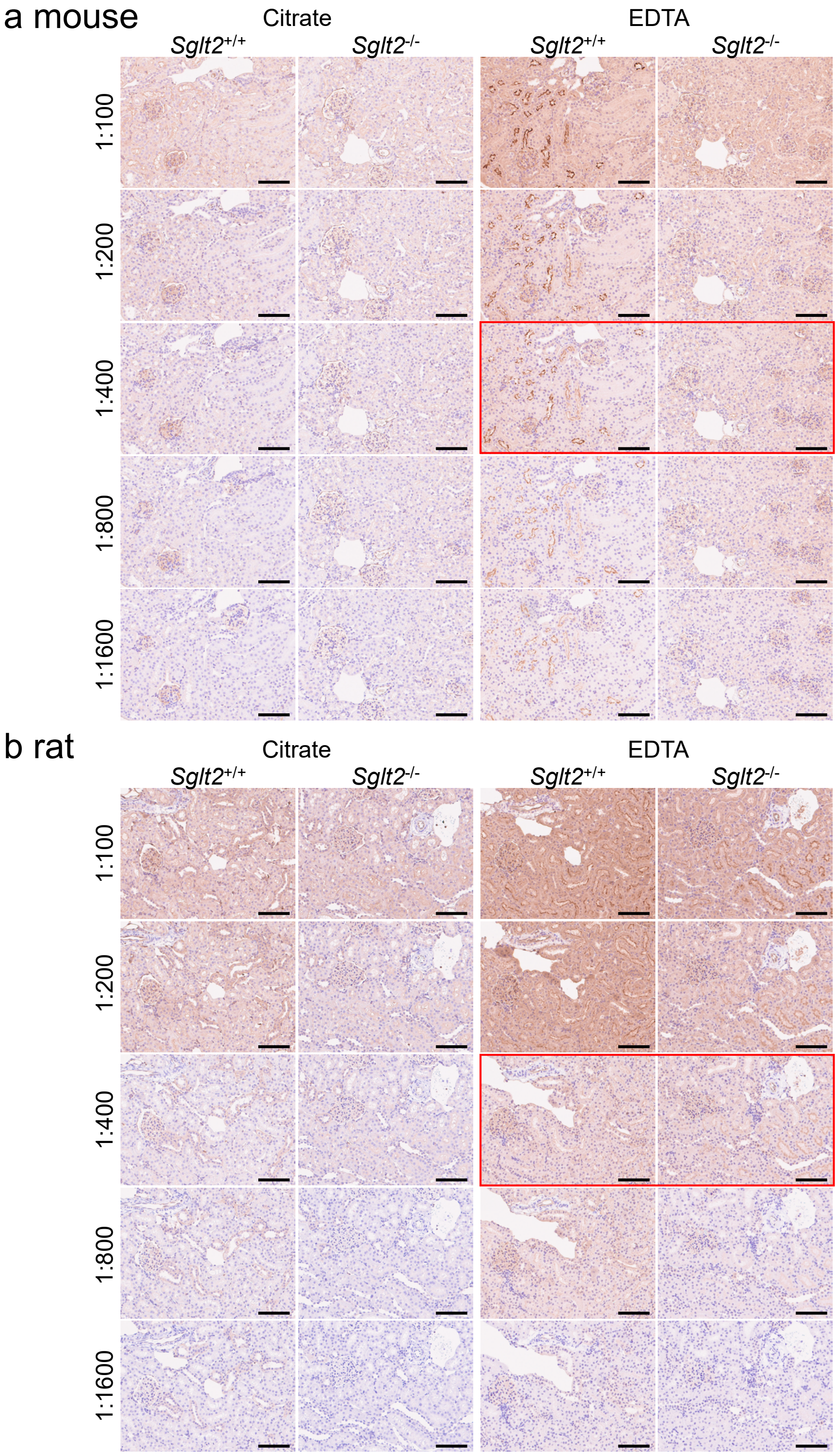


**Supplementary Figure S11. Optimization of anti-SGLT2 (sc-393350) antibody concentration and antigen retrieval in rodent kidney sections.**
Chromogenic detection of anti-SGLT2 (sc-393350) antibody immunostaining in the cortical regions of kidneys from wild-type (*Sglt2*$^{+/+}$) and *Sglt2*-deficient (*Sglt2*$^{-/-}$) mice (a) and rats (b). Five antibody dilutions (1:100 to 1:1600, twofold serial dilutions) and two antigen retrieval solutions (citrate buffer, pH 6.0 and EDTA buffer, pH 9.0) were tested. Nuclei were counterstained with hematoxylin. Images were captured using NanoZoomer-SQ slide scanner and exported to JPG file at x20 magnification using NDP.view2 software. The red box indicates the photo shown in Figure 1. Scale bar = 100 µm.

**Supplementary Figure S12**

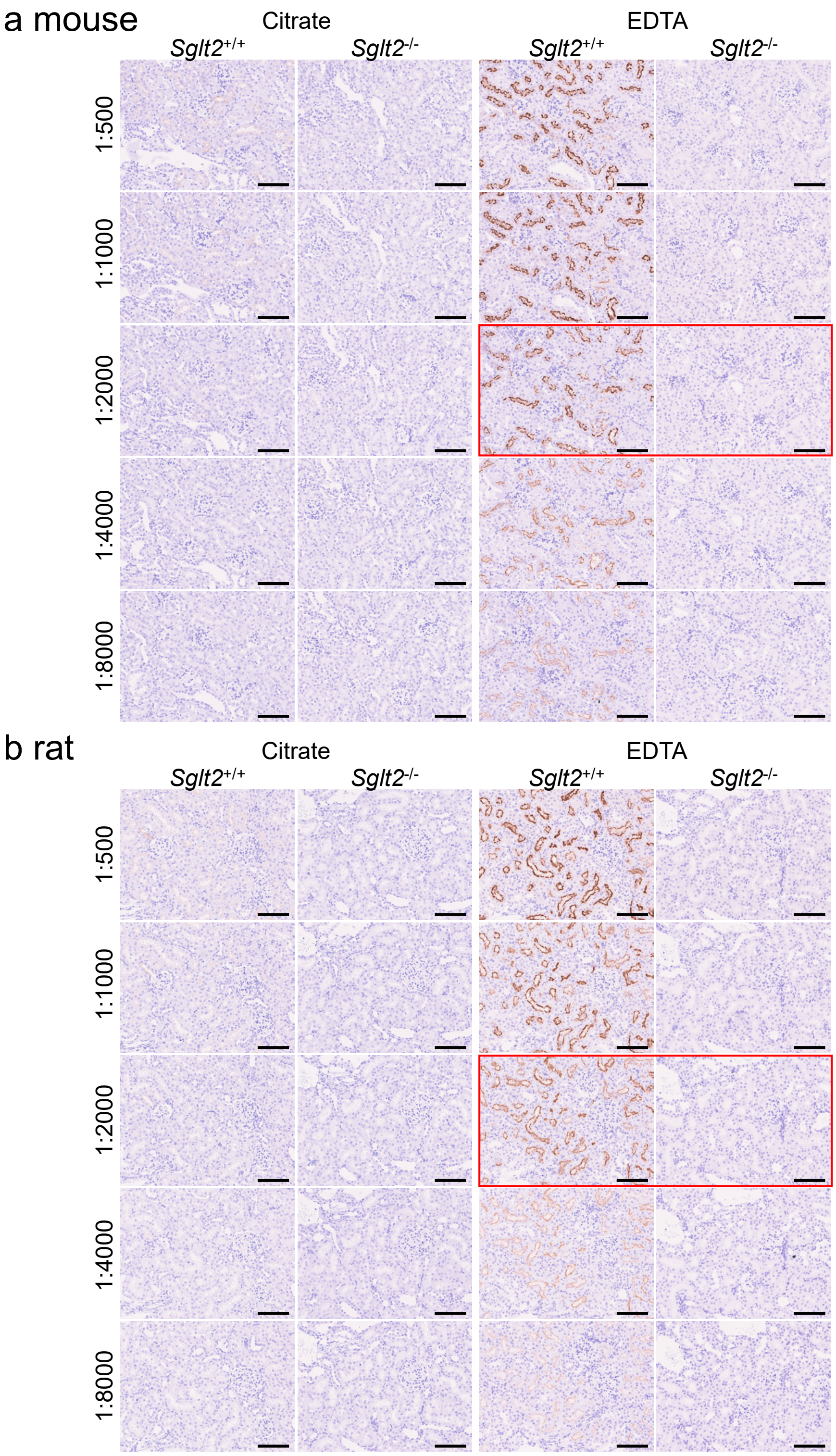


**Supplementary Figure S12. Optimization of anti-SGLT2 (HPA041603) antibody concentration and antigen retrieval in rodent kidney sections.**

Chromogenic detection of anti-SGLT2 (HPA041603) antibody immunostaining in the cortical regions of kidneys from wild-type (*Sglt2*$^{+/+}$) and *Sglt2*-deficient (*Sglt2*$^{-/-}$) mice (a) and rats (b). Five antibody dilutions (1:500 to 1:8000, twofold serial dilutions) and two antigen retrieval solutions (citrate buffer, pH 6.0 and EDTA buffer, pH 9.0) were tested. Nuclei were counterstained with hematoxylin. Images were captured using NanoZoomer-SQ slide scanner and exported to JPG file at x20 magnification using NDP.view2 software. The red box indicates the photo shown in Figure 1. Scale bar = 100 µm.

**Supplementary Figure S13**

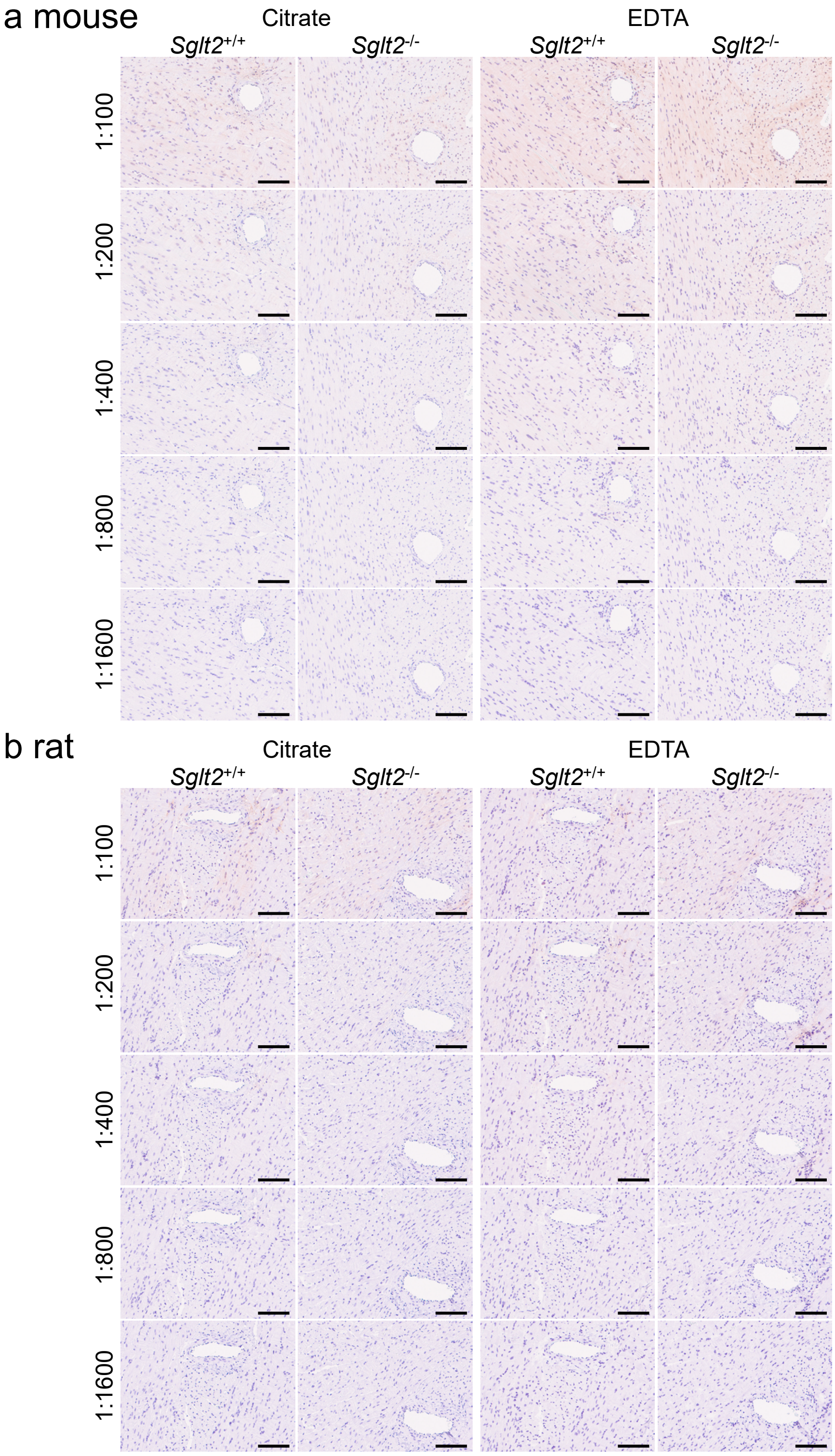


**Supplementary Figure S13. Immunostaining of anti-SGLT2 (ab37296) antibody in rodent heart sections.**
Immunostaining of anti-SGLT2 (ab37296) antibody in the left ventricular regions of hearts from wild-type (*Sglt2*$^{+/+}$) and *Sglt2*-deficient (*Sglt2*$^{-/-}$) mice (a) and rats (b). Five antibody dilutions (1:100 to 1:1600, twofold serial dilutions) and two antigen retrieval solutions (citrate buffer, pH 6.0 and EDTA buffer, pH 9.0) were tested. Nuclei were counterstained with hematoxylin. Images were captured using NanoZoomer-SQ slide scanner and exported to JPG file at x20 magnification using NDP.view2 software. Scale bar = 100 μm.

**Supplementary Figure S14**

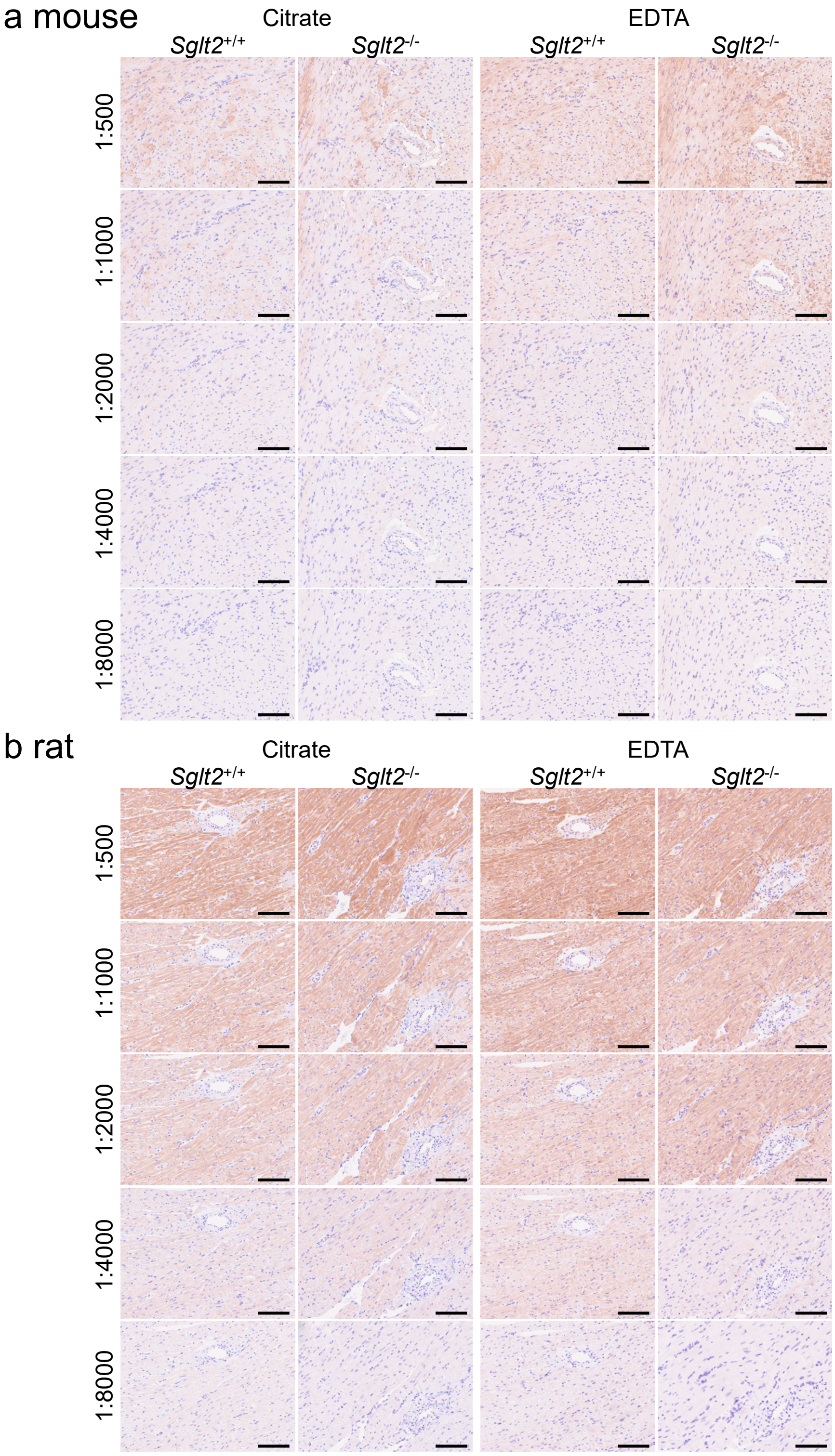


**Supplementary Figure S14. Immunostaining of anti-SGLT2 (ab85626) antibody in rodent heart sections.**

Immunostaining of anti-SGLT2 (ab85626) antibody in the left ventricular regions of hearts from wild-type (*Sglt2*$^{+/+}$) and *Sglt2*-deficient (*Sglt2*$^{-/-}$) mice (a) and rats (b). Five antibody dilutions (1:500 to 1:8000, twofold serial dilutions) and two antigen retrieval solutions (citrate buffer, pH 6.0 and EDTA buffer, pH 9.0) were tested. Nuclei were counterstained with hematoxylin. Images were captured using NanoZoomer-SQ slide scanner and exported to JPG file at x20 magnification using NDP.view2 software. Scale bar = 100 µm.

**Supplementary Figure S15**

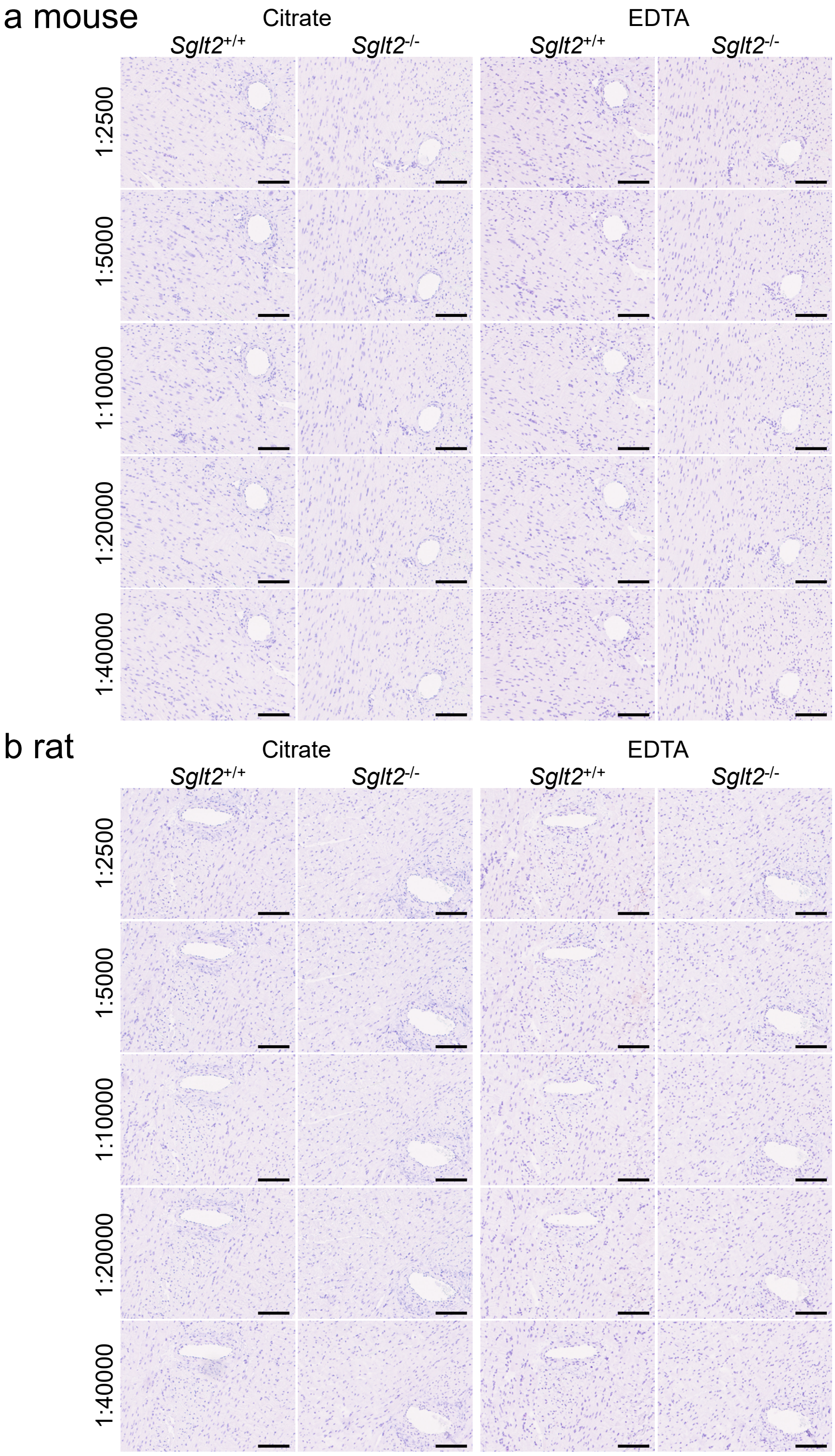


**Supplementary Figure S15. Immunostaining of anti-SGLT2 (ab306558) antibody in rodent heart sections.**
Immunostaining of anti-SGLT2 (ab306558) antibody in the left ventricular regions of hearts from wild-type (*Sglt2*$^{+/+}$) and *Sglt2*-deficient (*Sglt2*$^{-/-}$) mice (a) and rats (b). Five antibody dilutions (1:2500 to 1:5000, twofold serial dilutions) and two antigen retrieval solutions (citrate buffer, pH 6.0 and EDTA buffer, pH 9.0) were tested. Nuclei were counterstained with hematoxylin. Images were captured using NanoZoomer-SQ slide scanner and exported to JPG file at x20 magnification using NDP.view2 software. Scale bar = 100 µm.

**Supplementary Figure S16**

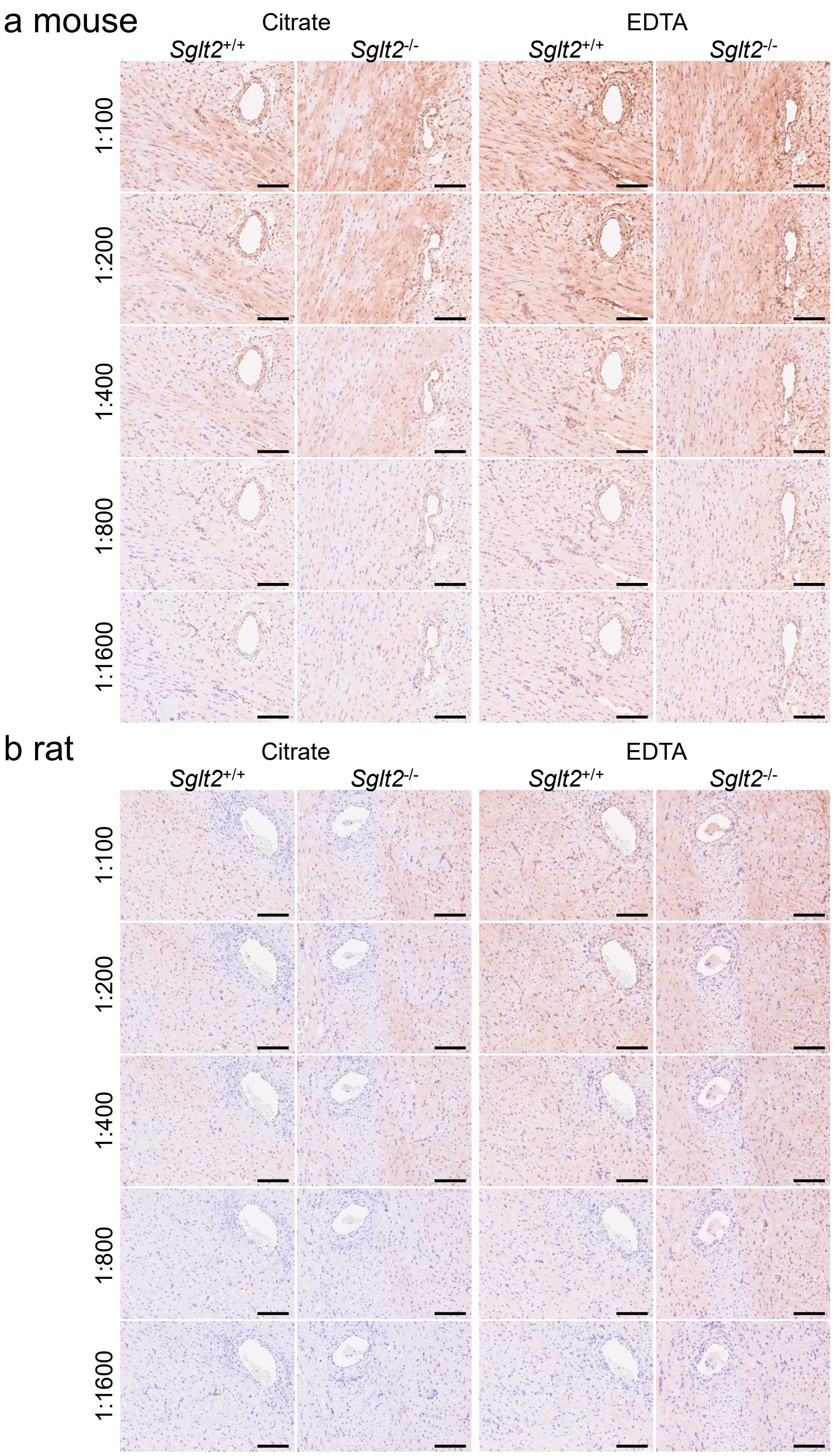


**Supplementary Figure S16. Immunostaining of anti-SGLT2 (20802) antibody in rodent heart sections.**

Immunostaining of anti-SGLT2 (20802) antibody in the left ventricular regions of hearts from wild-type (*Sglt2*$^{+/+}$) and *Sglt2*-deficient (*Sglt2*$^{-/-}$) mice (a) and rats (b). Five antibody dilutions (1:100 to 1:1600, twofold serial dilutions) and two antigen retrieval solutions (citrate buffer, pH 6.0 and EDTA buffer, pH 9.0) were tested. Nuclei were counterstained with hematoxylin. Images were captured using NanoZoomer-SQ slide scanner and exported to JPG file at x20 magnification using NDP.view2 software. Scale bar = 100 μm.

**Supplementary Figure S17**

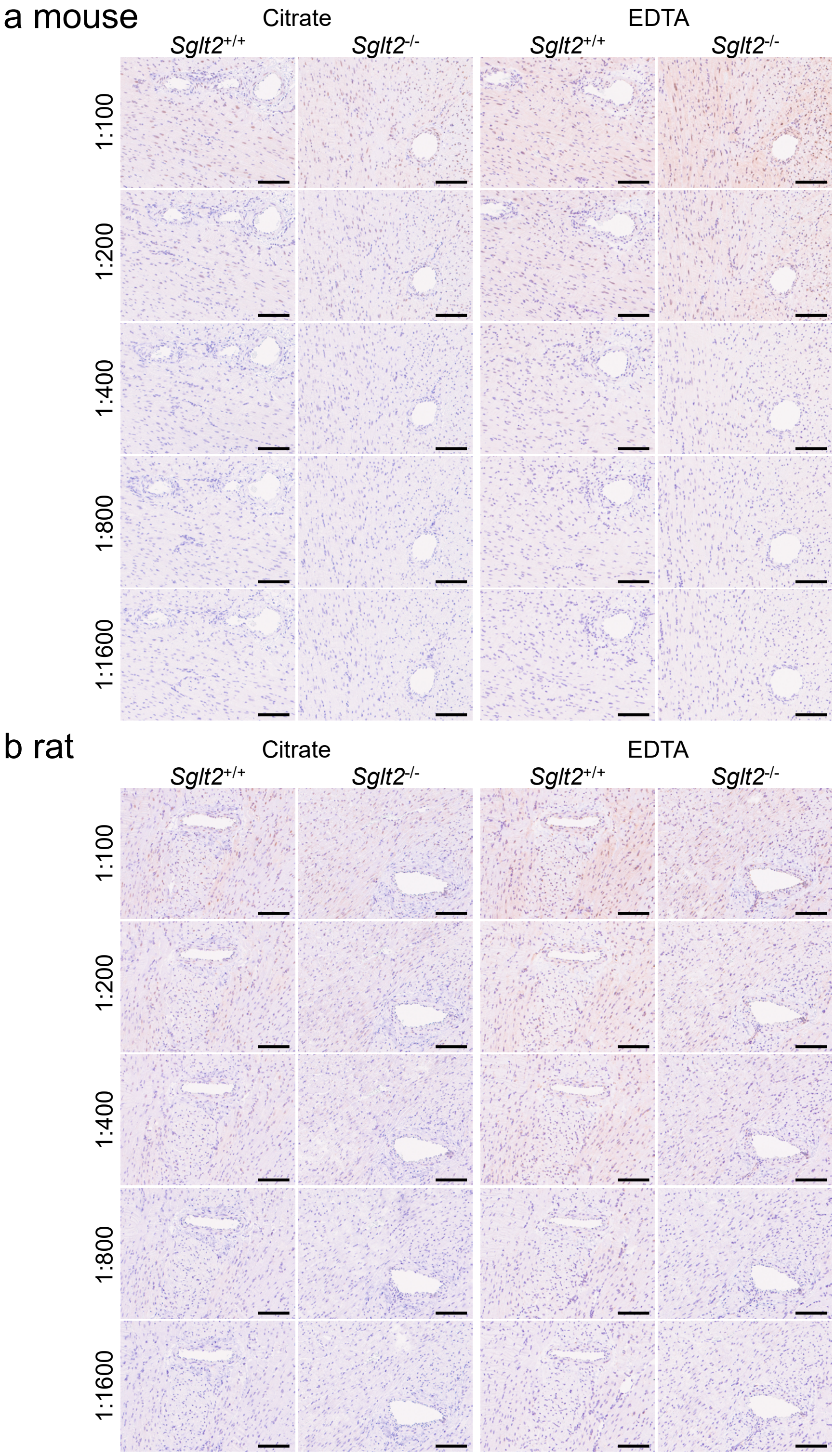


**Supplementary Figure S17. Immunostaining of anti-SGLT2 (14210) antibody in rodent heart sections.**

Immunostaining of anti-SGLT2 (14210) antibody in the left ventricular regions of hearts from wild-type (*Sglt2*$^{+/+}$) and *Sglt2*-deficient (*Sglt2*$^{-/-}$) mice (a) and rats (b). Five antibody dilutions (1:100 to 1:1600, twofold serial dilutions) and two antigen retrieval solutions (citrate buffer, pH 6.0 and EDTA buffer, pH 9.0) were tested. Nuclei were counterstained with hematoxylin. Images were captured using NanoZoomer-SQ slide scanner and exported to JPG file at x20 magnification using NDP.view2 software. Scale bar = 100 µm.

**Supplementary Figure S18**

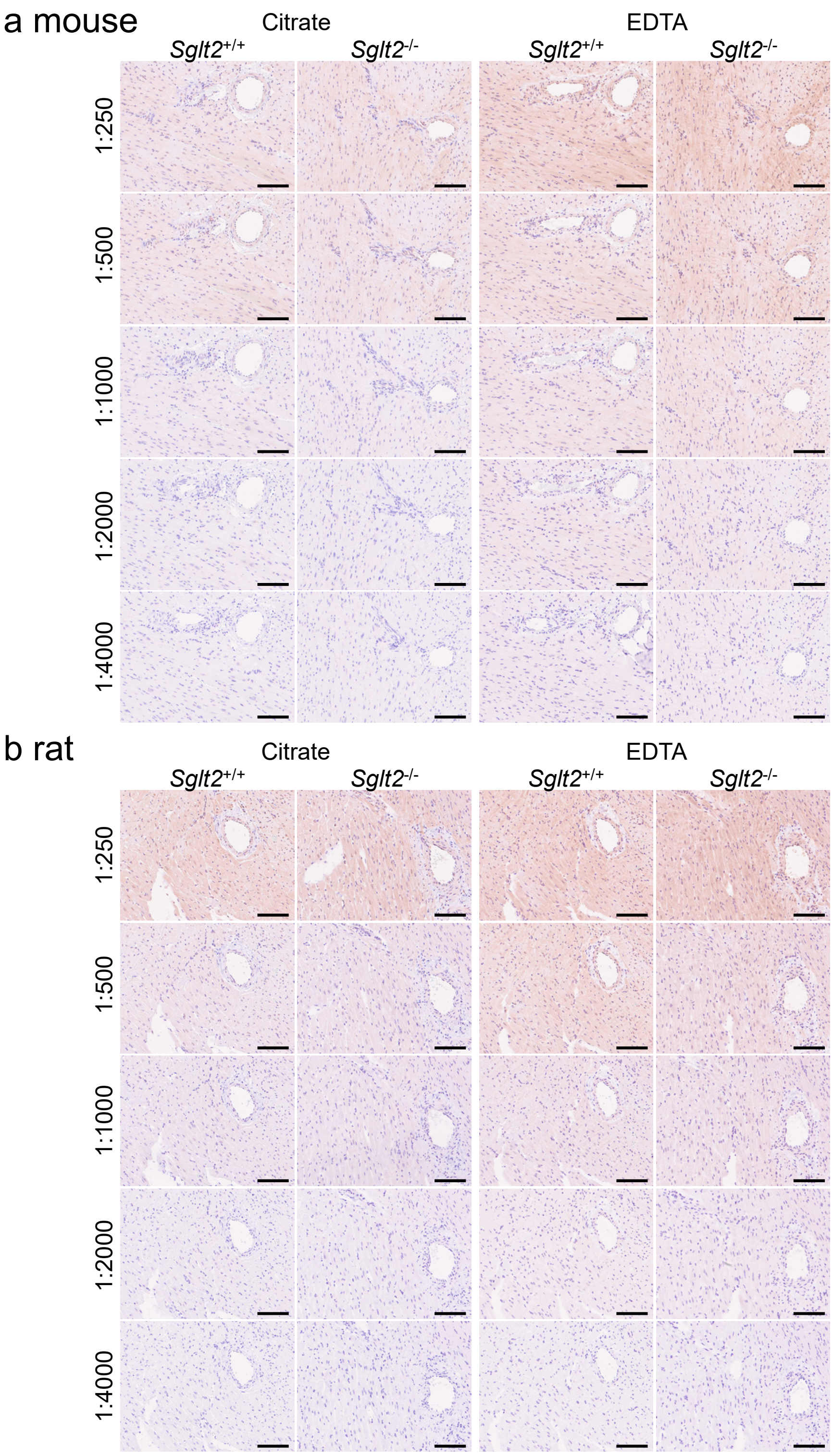


**Supplementary Figure S18. Immunostaining of anti-SGLT2 (24654-1-AP) antibody in rodent heart sections.**

Immunostaining of anti-SGLT2 (24654-1-AP) antibody in the left ventricular regions of hearts from wild-type (*Sglt2*$^{+/+}$) and *Sglt2*-deficient (*Sglt2*$^{-/-}$) mice (a) and rats (b). Five antibody dilutions (1:250 to 1:4000, twofold serial dilutions) and two antigen retrieval solutions (citrate buffer, pH 6.0 and EDTA buffer, pH 9.0) were tested. Nuclei were counterstained with hematoxylin. Images were captured using NanoZoomer-SQ slide scanner and exported to JPG file at x20 magnification using NDP.view2 software. Scale bar = 100 μm.

**Supplementary Figure S19**

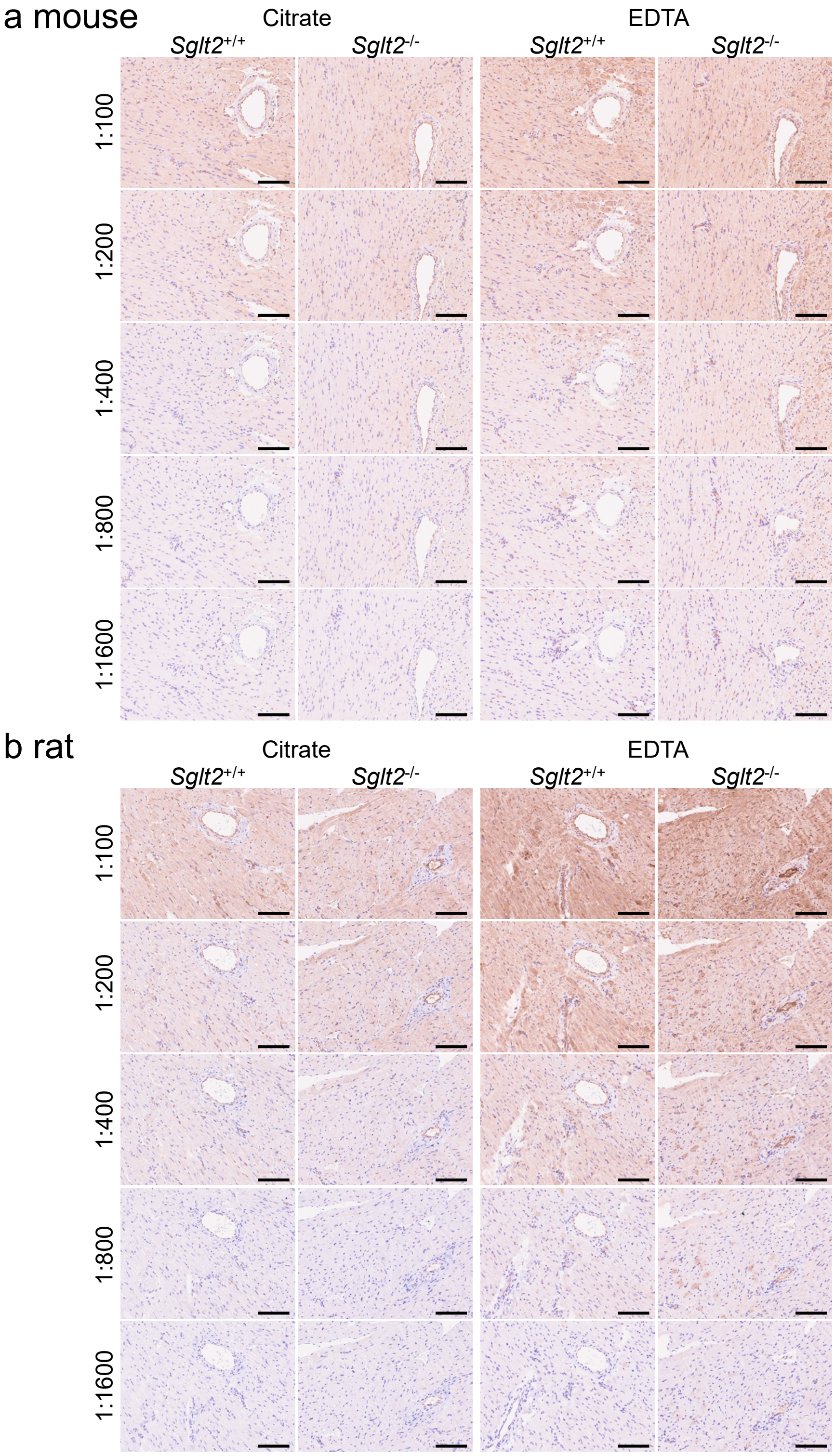


**Supplementary Figure S19. Immunostaining of anti-SGLT2 (sc-393350) antibody in rodent heart sections.**

Immunostaining of anti-SGLT2 (sc-393350) antibody in the left ventricular regions of hearts from wild-type (*Sglt2*$^{+/+}$) and *Sglt2*-deficient (*Sglt2*$^{-/-}$) mice (a) and rats (b). Five antibody dilutions (1:100 to 1:1600, twofold serial dilutions) and two antigen retrieval solutions (citrate buffer, pH 6.0 and EDTA buffer, pH 9.0) were tested. Nuclei were counterstained with hematoxylin. Images were captured using NanoZoomer-SQ slide scanner and exported to JPG file at x20 magnification using NDP.view2 software. Scale bar = 100 µm.

**Supplementary Figure S20**

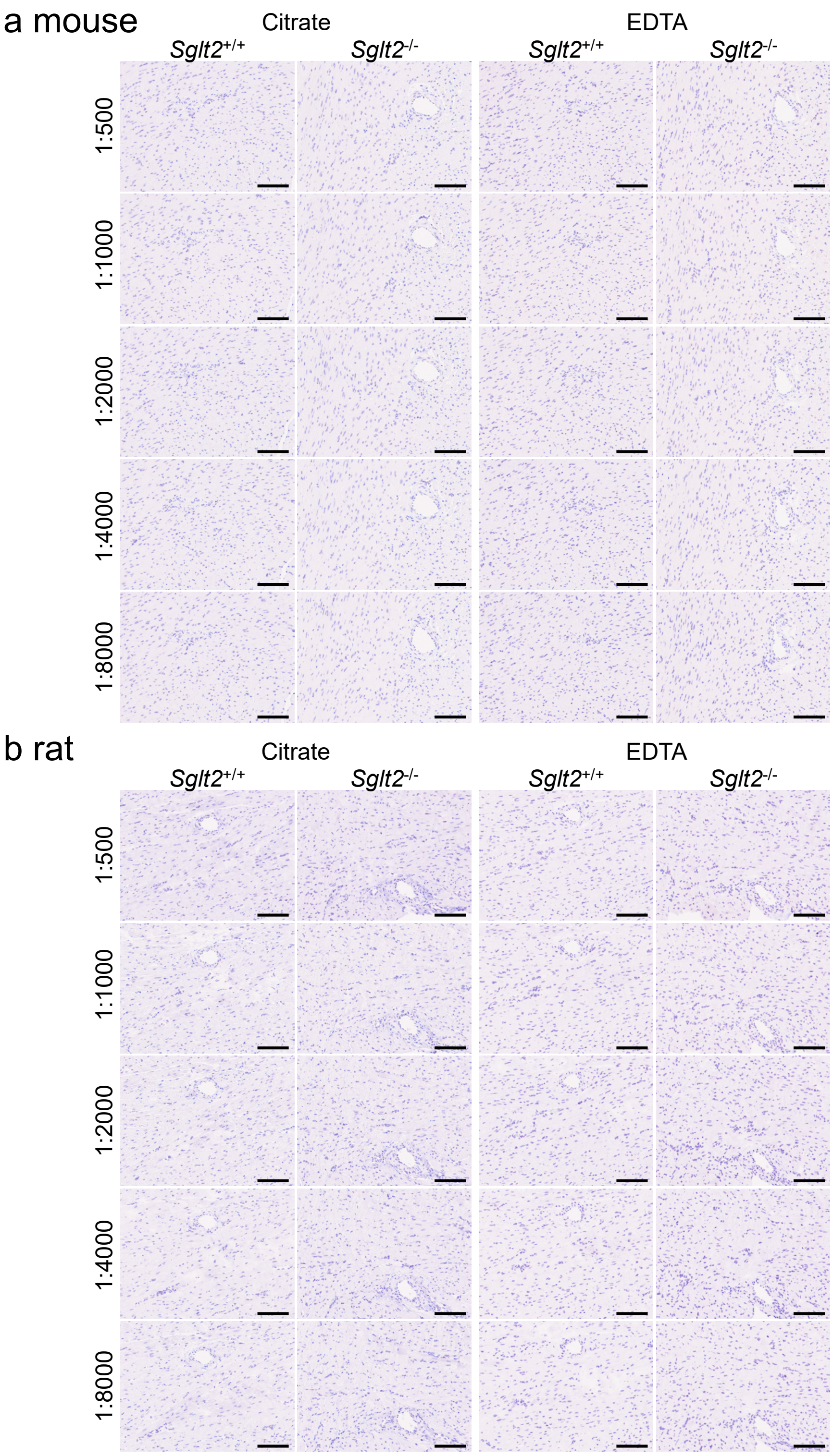


**Supplementary Figure S20. Immunostaining of anti-SGLT2 (HPA041603) antibody in rodent heart sections.**

Immunostaining of anti-SGLT2 (HPA041603) antibody in the left ventricular regions of hearts from wild-type (*Sglt2*$^{+/+}$) and *Sglt2*-deficient (*Sglt2*$^{-/-}$) mice (a) and rats (b). Five antibody dilutions (1:500 to 1:8000, twofold serial dilutions) and two antigen retrieval solutions (citrate buffer, pH 6.0 and EDTA buffer, pH 9.0) were tested. Nuclei were counterstained with hematoxylin. Images were captured using NanoZoomer-SQ slide scanner and exported to JPG file at x20 magnification using NDP.view2 software. Scale bar = 100 µm.

**Supplementary Figure S21**

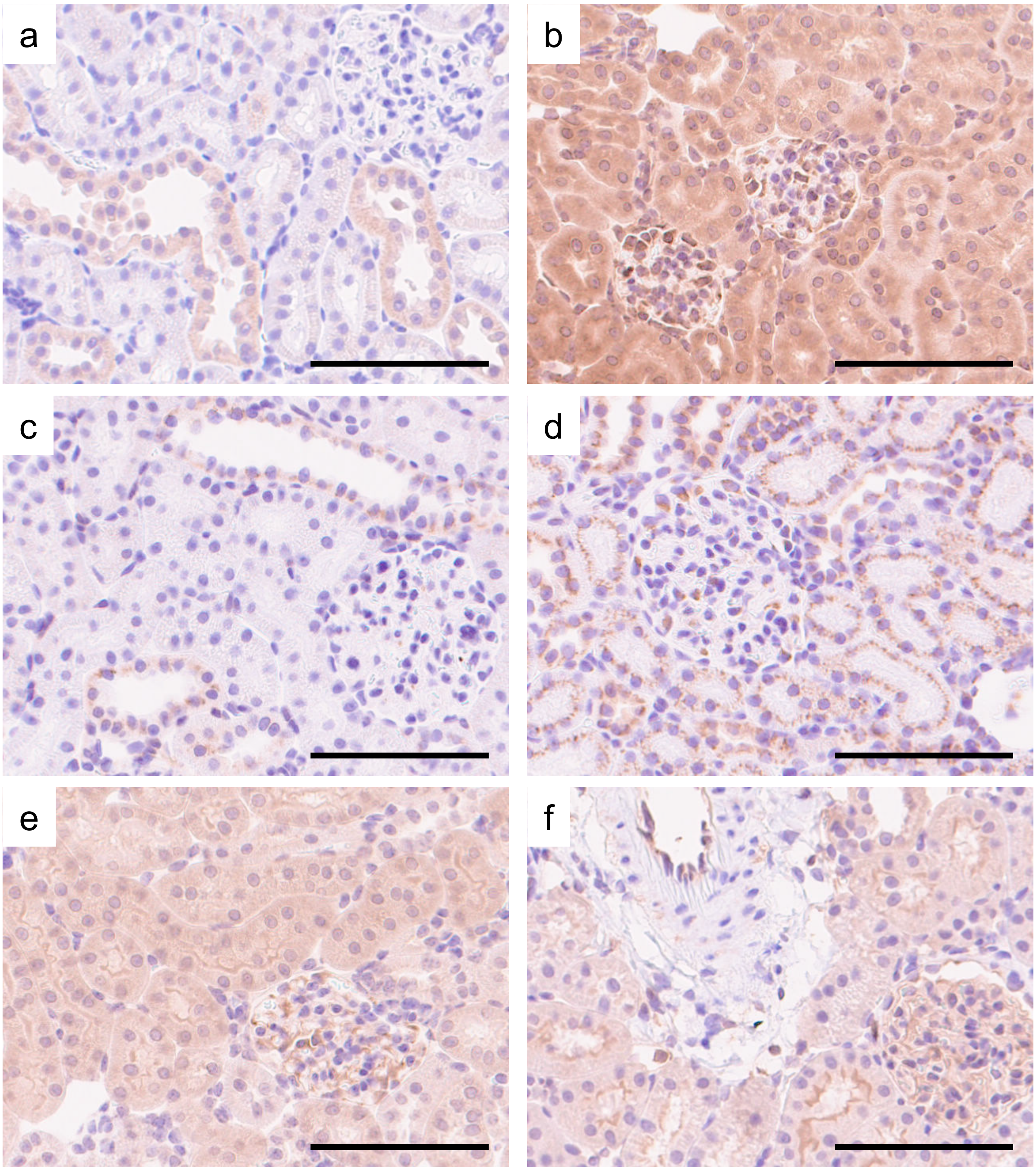


**Supplementary Figure S21. Non-specific staining of anti-SGLT2 antibodies in kidneys of *Sglt2*-deficient rodents.**

Representative images of non-specific immunostaining in (a) ab85626 (1:4000, citrate antigen retrieval, *Sglt2*-deficient rat), (b) 20802 (1:800, EDTA antigen retrieval, *Sglt2*-deficient mouse), (c) 20802 (1:800, citrate antigen retrieval, *Sglt2*-deficient rat), (d) 14210 (1:200, EDTA antigen retrieval, *Sglt2*-deficient rat), (e) 24654-1-AP (1:1000, EDTA antigen retrieval, *Sglt2*-deficient mouse), and (f) sc-393350 (1:400, EDTA antigen retrieval, *Sglt2*-deficient rat). Detailed staining patterns are described in Supplementary Table S3. Nuclei were counterstained with hematoxylin. Images were captured using NanoZoomer-SQ slide scanner and exported to JPG file at x20 magnification using NDP.view2 software. Scale bar = 100 μm.

**Supplementary Figure S22**

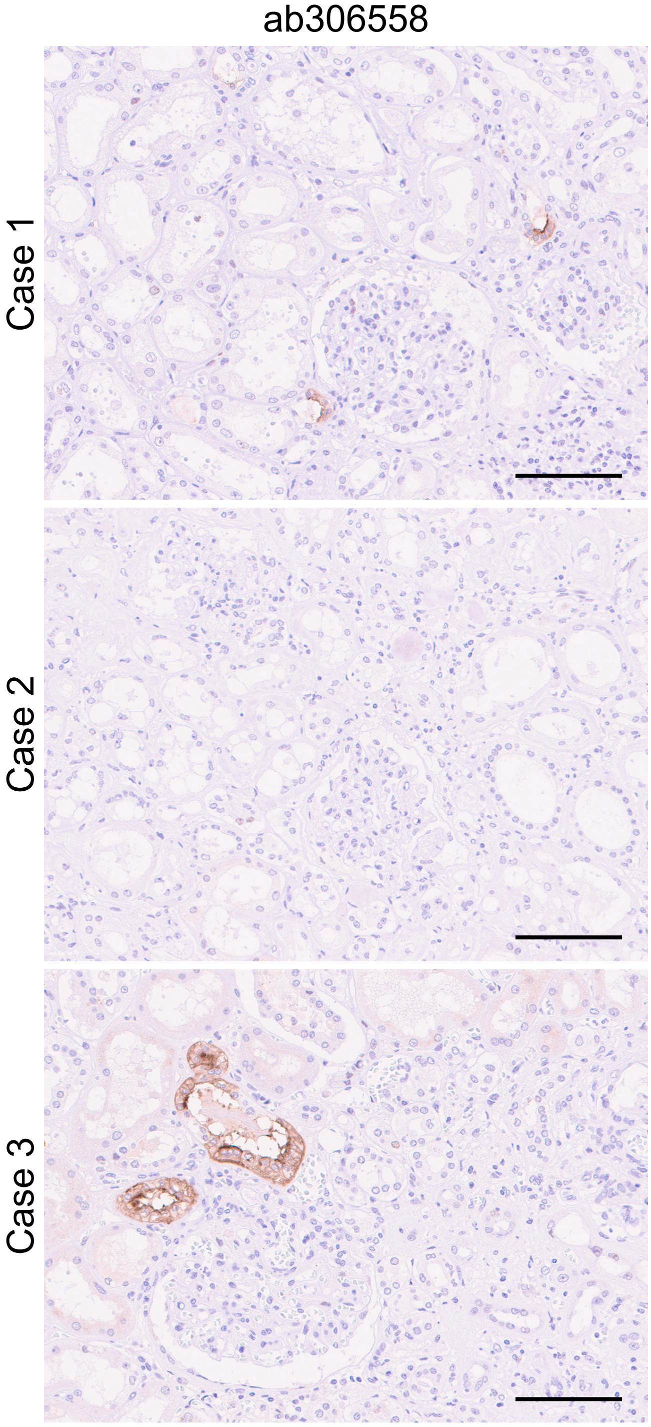


**Supplementary Figure S22. Immunostaining of anti-SGLT2 (ab306558) antibody in human autopsy kidney sections.**
Representative images of anti-SGLT2 (ab306558) antibody immunostaining in human autopsy kidney sections (n = 3). The antibody was diluted at 1:5000, with antigen retrieval by EDTA buffer (pH 9.0). Nuclei were counterstained with hematoxylin. Images were captured using NanoZoomer-SQ slide scanner and exported to JPG file at x20 magnification using NDP.view2 software. Scale bar = 100 µm.

**Supplementary Figure S23**

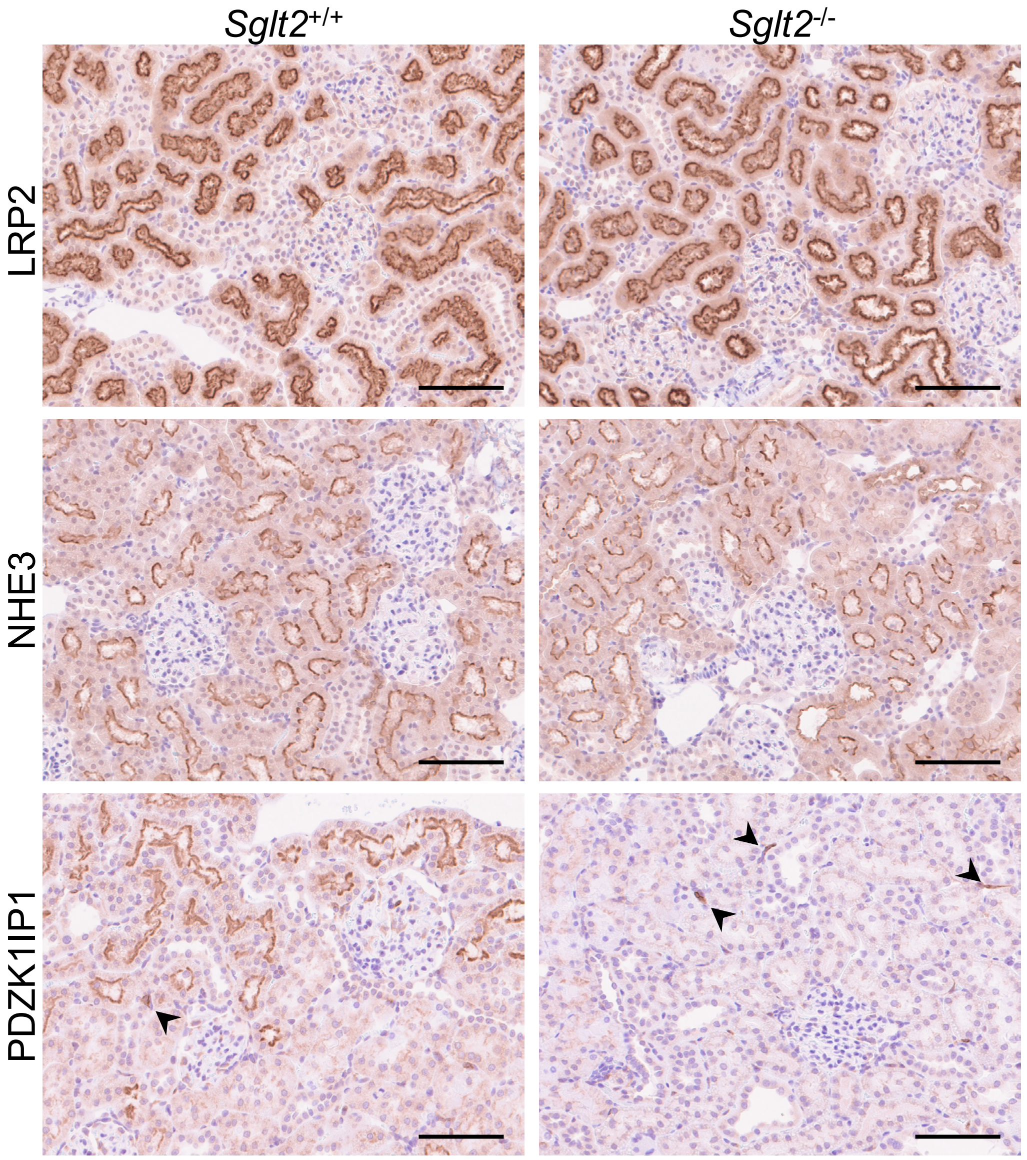


**Supplementary Figure S23. Immunostaining of proximal tubular markers in wild-type and *Sglt2*-deficient rodent kidneys.**
Chromogenic detection of LRP2 (megalin), NHE3 (SLC9A3), and PDZK1IP1 (MAP17) immunostaining in the cortical regions of kidneys from wild-type (*Sglt2*$^{+/+}$) and *Sglt2*-deficient (*Sglt2*$^{-/-}$) rats. Nuclei were counterstained with hematoxylin. Arrowheads indicate PDZK1IP1-positive infiltering blood cells. Images were captured using NanoZoomer-SQ slide scanner and exported to JPG file at x20 magnification using NDP.view2 software. Scale bar = 100 µm.

### Supplementary Figure S24

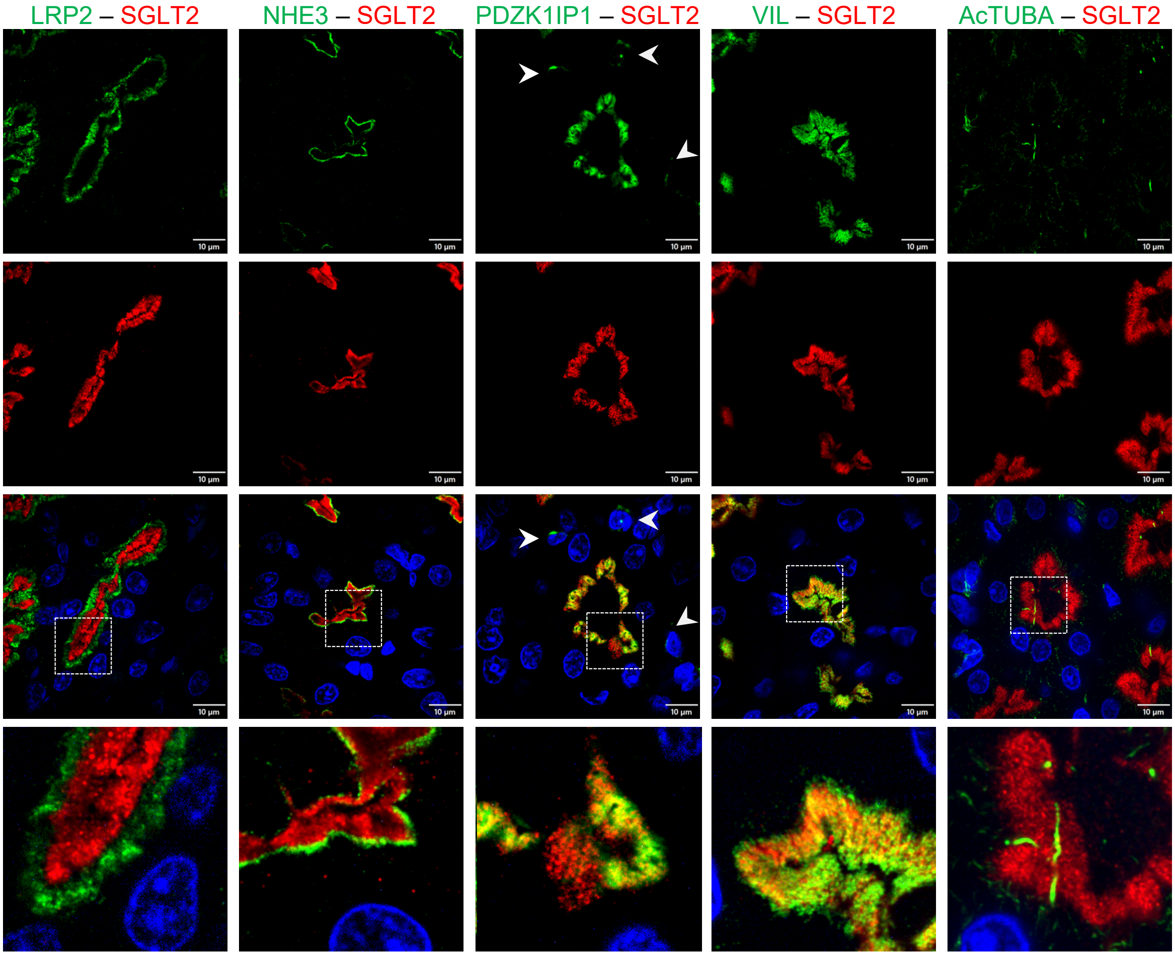


**Supplementary Figure S24. Double immunofluorescence staining of SGLT2 (ab306558) with proximal tubular markers in wild-type rat kidneys.**
Representative double-immunofluorescence images of SGLT2 (ab306558, red) with proximal tubular markers (green) in wild-type rat kidneys. Nuclei are counterstained with Hoechst 33342 (blue). Boxed regions of the merged images are magnified in the bottom panels. Arrowheads indicate PDZK1IP1-positive infiltering blood cells. Scale bar = 10 µm.

**Supplementary Figure S25**

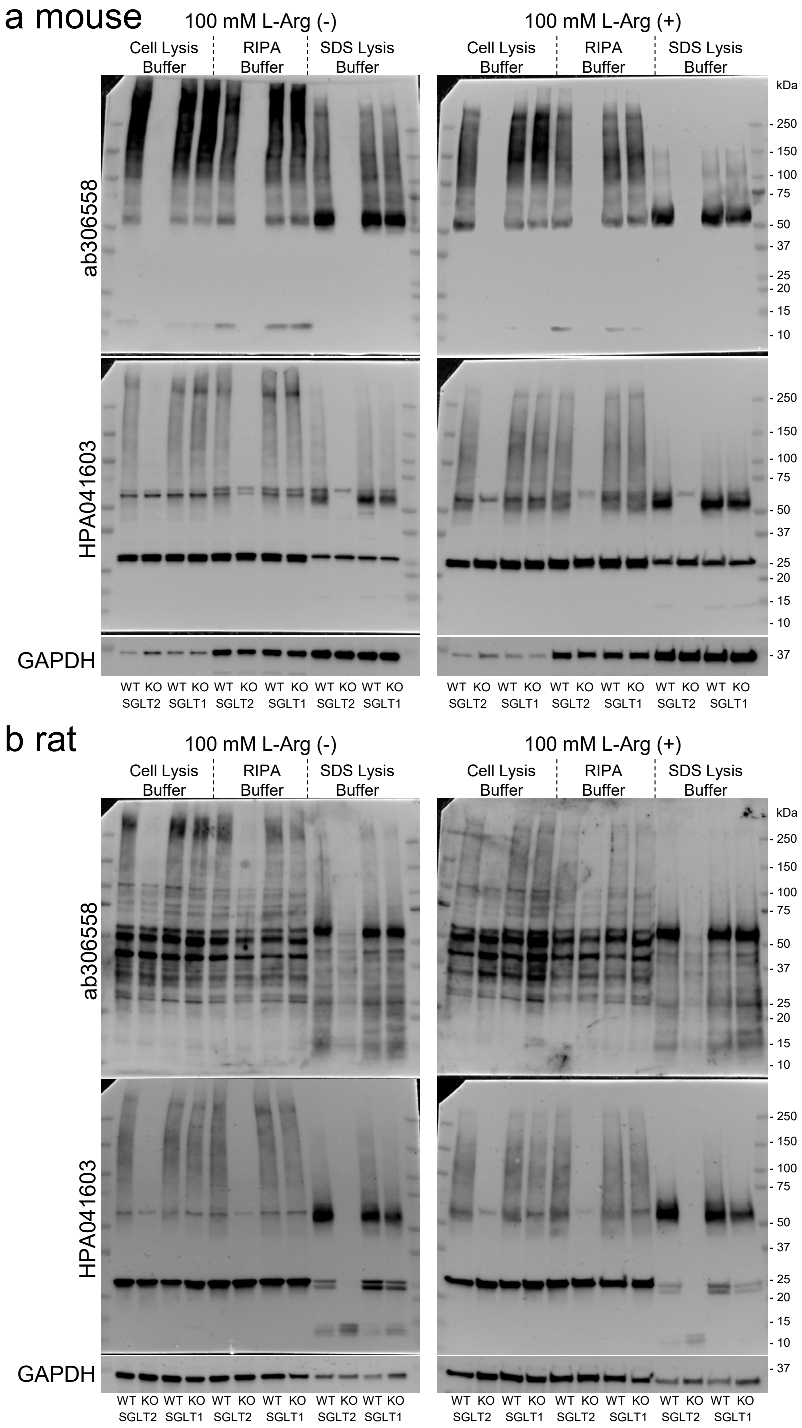


**Supplementary Figure S25. Effect of lysis buffer on SGLT2 band migration.**
(a, b) Representative full Western blot images of whole kidney lysate from *Sglt2*-deficient mouse, *Sglt1*-deficient mouse, and their wild-type littermates (a), and kidney cortex lysates from *Sglt2*-deficient rats, *Sglt1*-deficient rats, and their wild-type littermates (b). Lysates were prepared using three different lysis buffers: Cell Lysis Buffer, radioimmunoprecipitation assay (RIPA) buffer, or sodium dodecyl sulfate (SDS) lysis buffer. Twenty micrograms of total protein, with and without 100 mM L-arginine (L-Arg), were heat-denatured and separated electrophoresis. SGLT2 was detected with anti-SGLT2 antibodies, ab306558 and HPA041603. Glyceraldehyde 3-phosphate dehydrogenase (GAPDH) was used as a loading control. Molecular masses based on the protein ladder are indicated on the right side of the blots. WT, wild-type; KO, knockout.

**Supplementary Table S1:** Characteristics of 17 renal cell carcinoma (RCC) patients.

| Patient no. | Sex | Type | Stage (UICC 8th) | Grade* | Lymphovascular Invasion | Variant |
|---|---|---|---|---|---|---|
| 1 | F | Chromophobe RCC | pT1a | G2 | - | |
| 2 | F | papillary RCC | pT1a | G1 | - | |
| 3 | M | ccRCC | pT3a | G4 | + | sarcomatoid |
| 4 | M | ccRCC | pT3a | G4 | + | sarcomatoid |
| 5 | M | ccRCC | pT3a | G1 | - | |
| 6 | M | ccRCC | pT1b | G2 | - | |
| 7 | M | ccRCC | pT1a | G2 | - | |
| 8 | F | ccRCC | pT3a | G4 | - | sarcomatoid/rhabdoid |
| 9 | M | ccRCC | pT1a | G2 | - | |
| 10 | M | ccRCC | pT1a | G2 | - | |
| 11 | F | ccRCC | pT1a | G2 | - | |
| 12 | F | ccRCC | pT1a | G1 | - | |
| 13 | M | ccRCC | pT1b | G2 | + | |
| 14 | M | ccRCC | pT3a | G3 | + | |
| 15 | M | ccRCC | pT3a | G3 | + | |
| 16 | F | Chromophobe RCC | pT1a | G2 | - | |
| 17 | M | ccRCC | pT1a | G2 | - | |

UICC, Union for Intarnational Cancer Control; F, Female; M, Male; ccRCC, clear cell renal cell carcinoma.

* Fuhrman SA, Lasky LC, Limas C. Prognostic significance of morphologic parameters in renal cell carcinoma. Am J Surg Pathol 1982; 6:655-663.

**Supplementary Table S2:** Antibody information.

| Antigen | Vendor | Catalog no. | Clone no. | Host | IHC/IF | WB |
|---|---|---|---|---|---|---|
| AcTUBA | Sigma-Aldrich | T7451 | 6-11B-1 | mouse | 1:1000 | |
| GAPDH | Cell Signaling | 2118S | 14C10 | rabbit | | 1:5000 |
| LRP2 (megalin) | Santa Cruz | sc-515772 | H-10 | mouse | 1:1000 | |
| SLC9A3 (NHE3) | Millipore | MABN1813 | 3H3 | mouse | 1:500 | |
| PDZK1IP1 (MAP17) | Sigma-Aldrich | HPA014907 | - | rabbit | 1:1000 | |
| VIL | Santa Cruz | sc-58897 | 1D2C3 | mouse | 1:200 | |

IHC, immunohistochemistry; IF, immunofluorescence; WB, Western blotting; AcTUBA, acetylated alpha-Tubulin; GAPDH, glyceraldehyde-3-phosphate dehydrogenase; LRP2, low-density lipoprotein receptor-related protein 2; SLC9A3, solute carrier family 9 member A3; NHE3, sodium/hydrogen exchanger 3; PDZK1IP1, PDZ domain containing 1 interacting protein 1; MAP17, membrane-associated protein 17; VIL, villin.

**Supplementary Table S3:** Staining characteristics of anti-SGLT2 antibodies in kidneys of *Sglt2*-deficient rodents.

| Antibody | Mouse | Rat |
|---|---|---|
| ab37296 | Tubular cytoplasmic staining | Weak tubular cytoplasmic staining |
| ab85626 | Tubular cytoplasmic staining | Cytoplasmic staining in non-PTC tubules (a) |
| ab306558 | No staining | No staining |
| 20802 | Glomerular and tubular cytoplasmic staining (b) | Nuclear staining (citrate); vesicular-like staining in non-PTC tubules (EDTA, c) |
| 14210 | Nuclear staining (citrate); weak tubular cytoplasmic staining (EDTA) | Glomerular staining and vesicular-like staining in tubules (d) |
| 24654-1-AP | Glomerular and weak tubular cytoplasmic staining (e) | Weak tubular cytoplasmic staining |
| sc-393350 | Vascular endothelial cell staining and PTC brush border staining | Vascular endothelial cell staining and PTC brush border staining (f) |
| HPA041603 | No staining | No staining |

a–f: corresponding images are shown in Supplementary Figure S21.

Antigen retrieval buffer is indicated in parentheses when staining patterns differed between citrate and ethylenediaminetetraacetic acid (EDTA).

PTC, proximal tubular cell.
